# Proteome-scale inference of transcription factor binding motifs from predicted protein-DNA structures

**DOI:** 10.64898/2026.11.18.767614

**Authors:** Benjamin Iovino, Nash Bulaong, Louis Boahene, William Law, Nathan Hammond, Mark Potts, Rosio Alvarez, Joseph L. DeRisi, John Pak, Yang Joon Kim, Loïc A. Royer

## Abstract

Transcription factors (TFs) control development and cell identity by binding sequence-specific DNA motifs, yet the binding specificities of most TFs remain unknown because experimental binding assays do not scale. In zebrafish, only one-third of TFs have assigned motifs, nearly all inferred by homology from human or mouse experimental data. Here we present a proteome-scale computational screen that infers DNA-binding specificity from predicted protein–DNA structures. We use AlphaFold3 to model zebrafish TFs with candidate motif sequences derived from accessible chromatin, then use NA-MPNN to infer the DNA motifs from the most confident predictions. Across 210 TFs with experimentally characterized orthologs, predicted motifs match the known motif for 44%. Propagated through sequence clusters, these predictions form ZebraMotif, a motif database for all 2,543 zebrafish TFs. ZebraMotif covers 2.8 times as many TFs as CisBP’s zebrafish collection and links accessible regulatory elements to candidate upstream regulators genome-w ide. Because the approach requires only a proteome and a set of candidate motifs, it is applicable to any species lacking experimental binding data.

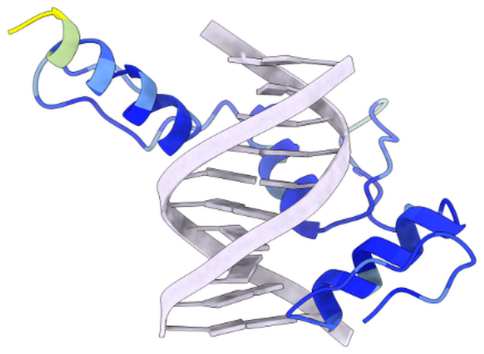

## Introduction

Transcription factors (TFs) are sequence-specific DNA-binding proteins that regulate gene expression by binding short motifs within *cis*-regulatory elements to activate or repress transcription.^1^ They control development, cell identity, and responses to the environment, and mutations in either TFs or their binding sites can cause human disease.^2, 3^ Defining the DNA-binding specificities of TFs, often represented as position weight matrices (PWMs) over a recognition sequence of 8–14 base pairs,^4^ is therefore necessary to reconstruct the gene regulatory networks that coordinate these processes. Binding motifs are determined experimentally by *in vitro* methods such as protein binding microarrays (PBMs),^5^ SELEX,^6^ SMiLE-seq,^7^ and more recently GHT-SELEX,^8^ as well as by *in vivo* chromatin immunoprecipitation followed by sequencing (ChIP-seq).^9^ These efforts have populated large curated databases, such as CisBP^10^ and JASPAR,^11^ but because the assays are hard to scale, coverage remains sparse outside of human and a few well-studied model organisms. Existing computational methods do not fill in the missing motifs: homology transfer is unreliable across large evolutionary distances,^12^ sequence-based models such as DeepBind, BPNet, and AlphaGenome learn from experimental binding or functional genomic data and do not directly infer motifs for TFs lacking such data,^13–15^ and recognition-code models predict approximate specificities for only a few families.^16, 17^

Structural studies of protein–DNA complexes have established that specificity is encoded in the chemical and geometric complementarity of the protein–DNA interface, with crystal structures revealing two recognition modes: direct readout, in which side chains form hydrogen bonds with specific nucleotides in the major groove, and indirect readout, in which proteins sense the local shape and electrostatic properties of the DNA itself, such as arginine insertion into narrow, AT-rich minor grooves.^18^ Deep learning models trained on large sequence and structural databases, including AlphaFold3 (AF3)^19^ and related models,^20–22^ can now predict protein–DNA complex structures accurately enough to capture interface geometry without experimental binding data, so specificity might be predicted directly from sequence. Structure-conditioned models such as DeepPBS^23^ and NA-MPNN^24^ can infer DNA base preferences from protein–DNA complexes, and the interface predicted template modeling (ipTM) score of predicted complexes can discriminate allele-specific binding preferences at known human TF binding sites.^25^ We therefore ask whether confidence-guided selection of predicted complexes, coupled to structureconditioned specificity inference, can recover TF motifs across diverse DNA-binding domain (DBD) families at proteome scale without experimental binding data for each TF.

Zebrafish (*Danio rerio*) is well-suited to test this. It is a widely used vertebrate model for dissecting developmental gene regulatory networks, with well-characterized embryogenesis, a fully sequenced genome, and emerging single-cell atlases that resolve cell-type-specific gene expression and chromatin accessibility.^26–29^ Despite this, experimentally derived DNA-binding specificities of zebrafish TFs are largely absent from public databases: of the 2,543 annotated TFs,^30^ CisBP assigns motifs to about one-third, nearly all by homology inference from human or mouse experiments.^10^

Here, we present a structure-based *in silico* screen that infers TF–DNA binding specificity at proteome scale, and apply it to zebrafish. Using AF3 and NA-MPNN, we screened over two million predicted TF-DNA structures and constructed ZebraMotif, a motif database covering 2,543 zebrafish TFs and browsable at zebramotif.apps.czbiohub.org. We show that structural confidence discriminates cognate from non-cognate DNA across diverse DBD families, and that across positive-control TFs with well-conserved JASPAR orthologs, predicted motifs match the experimental reference more often than motifs from the same DBD family. The same screen can be run on any species with an annotated genome and a source of candidate motifs to supply the motifs needed to reconstruct its gene regulatory networks.

## Results

### AF3 confidence separates cognate from non-cognate DNA for most transcription factors

Predicting a motif for every TF requires a signal that identifies when a TF is bound to its correct DNA sequence, without prior binding data for that TF. We therefore first tested whether AF3 distinguishes cognate from non-cognate DNA by co-folding each with the TF and comparing the confidence scores between them (Fig. 1A). We assembled a benchmark of 29 zebrafish TFs with direct experimental evidence in CisBP^10^ or a near-identical human ortholog in JASPAR^11^ (2024 CORE vertebrates, redundant; see Methods), together spanning 20 DBD families that differ in multiple-sequence-alignment (MSA) depth and percent identity to the structural templates returned from searching PDB (Supp. Fig. 1A). Predicted DBDs superpose closely on their PDB templates, but fold accuracy does not identify an accurate DNA-bound complex (Supp. Fig. 1B,C), so we turned to AF3’s confidence in the interface.

**Fig. 1.**
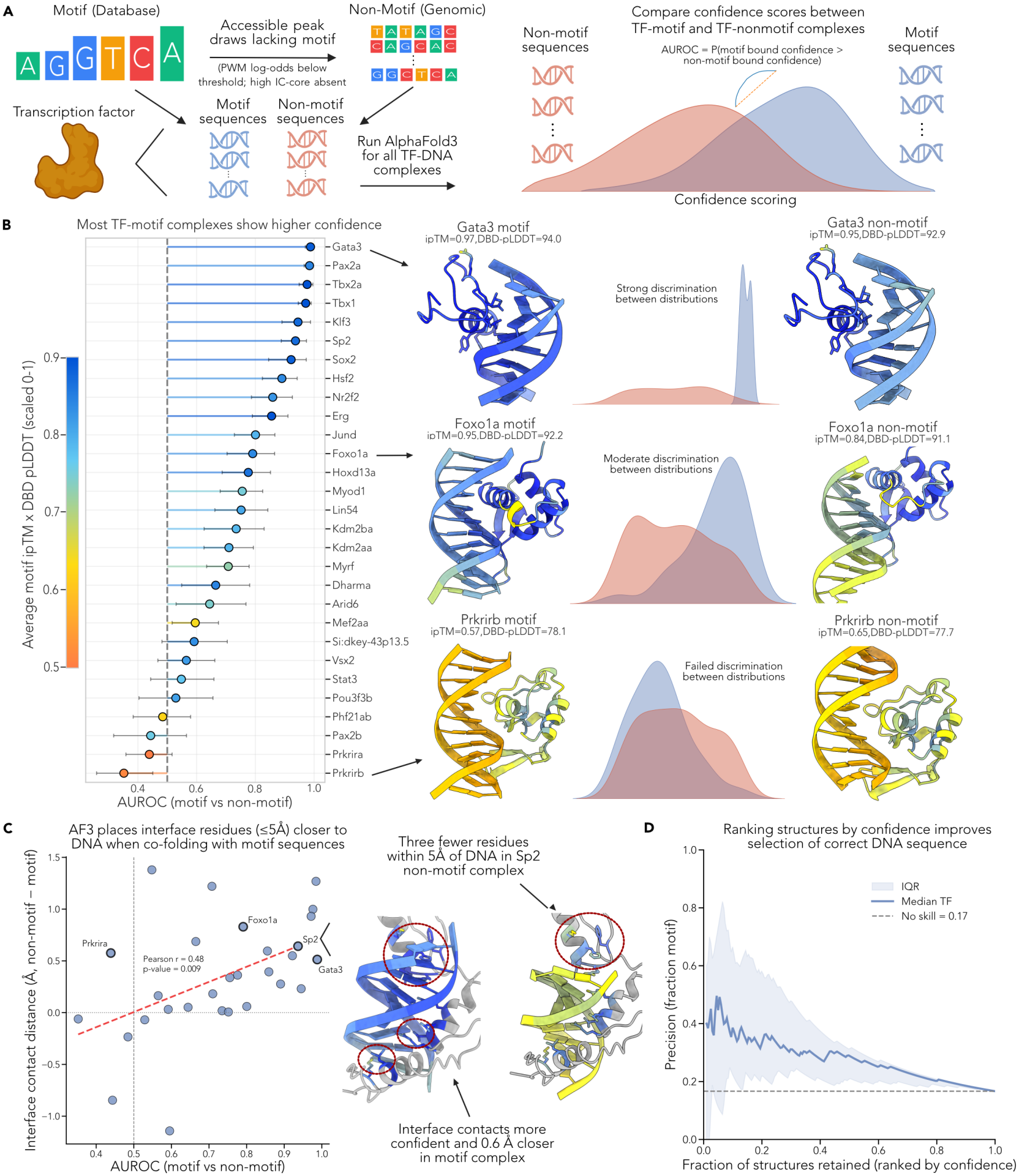
AlphaFold3 (AF3) confidence metrics can discriminate between motif and non-motif bound structures. **(A)** AF3 is benchmarked by co-folding each TF with motif and non-motif DNA. Each transcription factor (TF) is co-folded by AF3 with two sets of DNA: motif sequences sampled from its position weight matrix (PWM), and non-motif sequences drawn from accessible genomic peaks that lack the motif. All resulting protein–DNA complexes are scored by AF3 confidence. Discrimination is quantified per TF as the area under the receiver operating characteristic curve (AUROC), which is the probability that a motif-bound complex scores higher than a non-motif-bound complex. **(B)** AF3 confidence discriminates motif from non-motif DNA for most TFs. Per-TF AUROC (motif versus non-motif) for the 29 TFs, colored by confidence. Representative motif- and non-motif-bound structures are shown for strong (Gata3), moderate (Foxo1a) and failed (Prkrirb) discrimination, with ipTM and DNA-binding domain (DBD) pLDDT annotated and the confidence distributions inset. **(C)** Residues drawn closer to cognate DNA in the interface. Mean change in side-chain-to-nucleobase distance at interface residues (non-motif minus motif) versus each TF’s confidence AUROC from (B) (Pearson *r* = 0.48, *p* = 0.009; *n* = 29). Right, Sp2 motif- and non-motif-bound structures, with interface residues that lie closer to DNA in the motif complex circled. **(D)** Ranking structures by confidence improves selection of cognate DNA sequences. Precision (fraction of structures at the threshold that are motif-bound) versus the fraction of structures accepted when ranking by confidence, median across TFs (line), interquartile range (shaded) and the no-skill base rate (dashed; 30/180 = 0.17).

We co-folded each TF with 30 DNA sequences sampled from its cognate motif, enough to reconstruct the source PWM (Supp. Fig. 2), and 150 GC- and length-matched non-motif sequences from accessible-chromatin peaks (Fig. 1A; Methods). We scored each complex with a composite confidence, *S*_conf_, that combines ipTM with the predicted local distance difference test (pLDDT) averaged over the DBD (DBD-pLDDT; Methods). The DBD is the annotated domain that AF3 most confidently positions against the DNA (e.g. the paired box of Pax2a; Supp. Fig. 3; Methods). *S*_conf_ separates motif from non-motif complexes for most TFs, significantly for 20 of 29 (one-sided Mann–Whitney U, *p* < 0.05), with a mean per-TF area under the receiver operating characteristic curve (AUROC) of 0.73 and a mean average precision of 0.44 against a chance rate of 0.17 (Fig. 1B). Interface-only scores perform comparably (Supp. Fig. 4C), but DBD-pLDDT rescues a few cases that the interface-specific metrics do not, such as Mef2aa and Pou3f3b (Supp. Fig. 4D), so we use *S*_conf_ for all structure ranking after the first-pass motif filter.

**Fig. 2.**
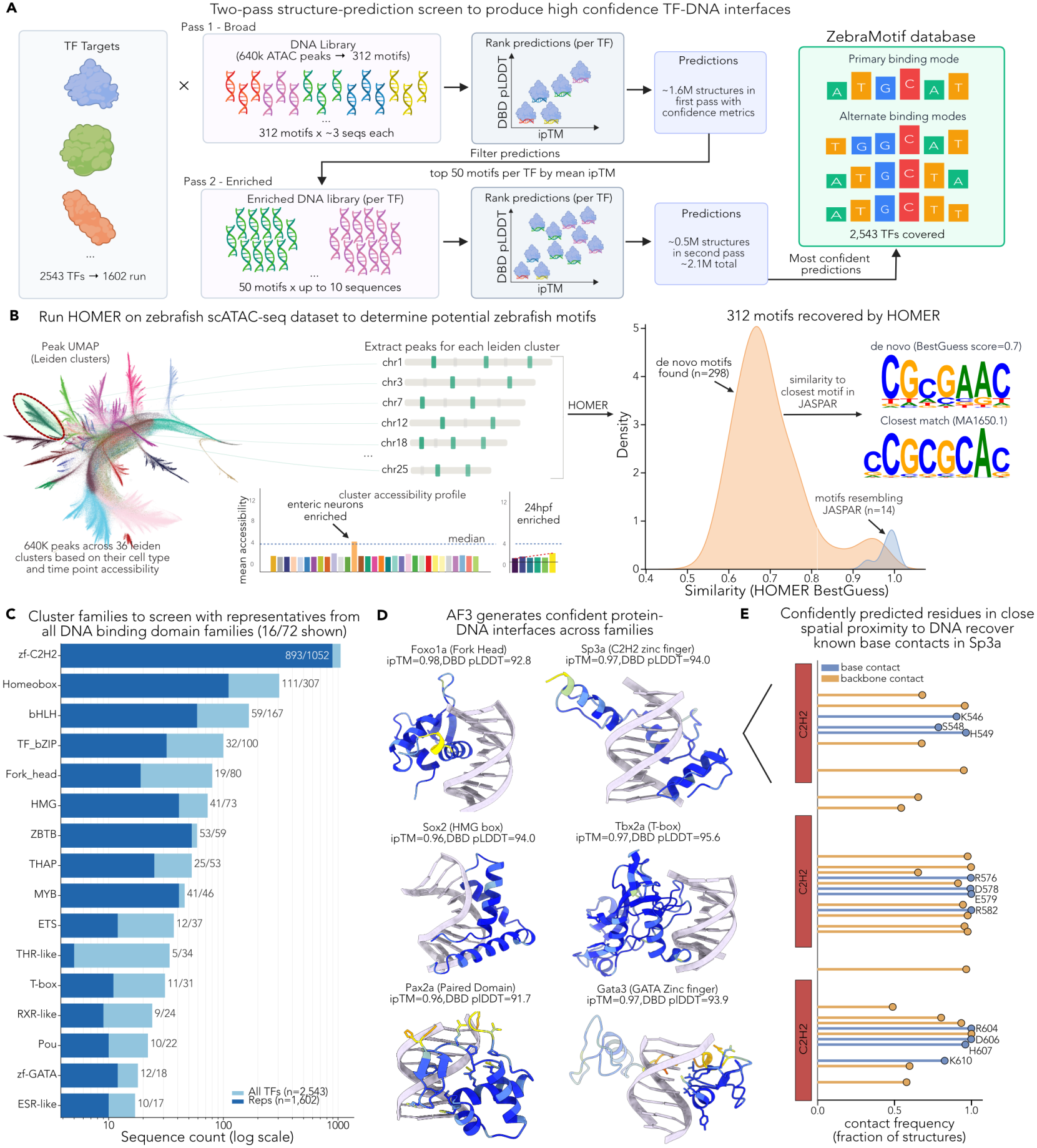
A proteome-scale, structure-guided screen predicts DNA-binding motifs for zebrafish transcription factors (TFs). **(A)** A two-pass screen searches the candidate-motif library against TF representatives. The 1,602 representatives are screened against the 312-motif library. A broad first pass (~1.6M predicted structures) identifies each target’s highest-confidence hit motifs. An enriched second pass samples the top 50 motifs per target more deeply (~0.5M structures). Confident hits from both passes feed per-TF position probability matrix (PPM) prediction, yielding the ZebraMotif database, covering 2,543 zebrafish TFs with primary and alternate binding modes. **(B)** The DNA library is built *de novo* from accessible chromatin in zebrafish. HOMER motif discovery within each of 36 Leiden clusters of the Zebrahub-Multiome snATAC-seq atlas (~640,000 peaks, 10–24 hpf) yields 312 non-redundant motifs. Density plot shows the BestGuess score (HOMER’s similarity to the closest JASPAR-vertebrates match) of each *de novo* motif. Example *de novo* motif (top, BestGuess score 0.7) and its closest JASPAR match (bottom, MA1650.1, ZBTB14). **(C)** TF targets are all annotated zebrafish TFs (AnimalTFDB4; n = 2,543), reduced to representatives (n = 1,602) by family-appropriate clustering. Sequence counts per DNA-binding domain (DBD) family (log scale; 16 of 72 families shown). **(D)** AlphaFold3 generates confident protein–DNA interfaces across DBD families. Representative predicted complexes for Foxo1a (Fork Head), Sp3a (C2H2 zinc finger), Sox2 (HMG box), Tbx2a (T-box), Pax2a (Paired domain), and Gata3 (GATA zinc finger). Proteins colored by pLDDT, DNA in grey, with ipTM and DBD-pLDDT annotated. **(E)** Recovered contacts reproduce known base-reading residues. For Sp3a, per-residue frequency (fraction of ensemble structures) of residues within 5 Å of DNA, split into base-directed (blue) and backbone (orange) contacts across its three C2H2 fingers. Base-directed residues coincide with the canonical recognition-code positions.

**Fig. 3.**
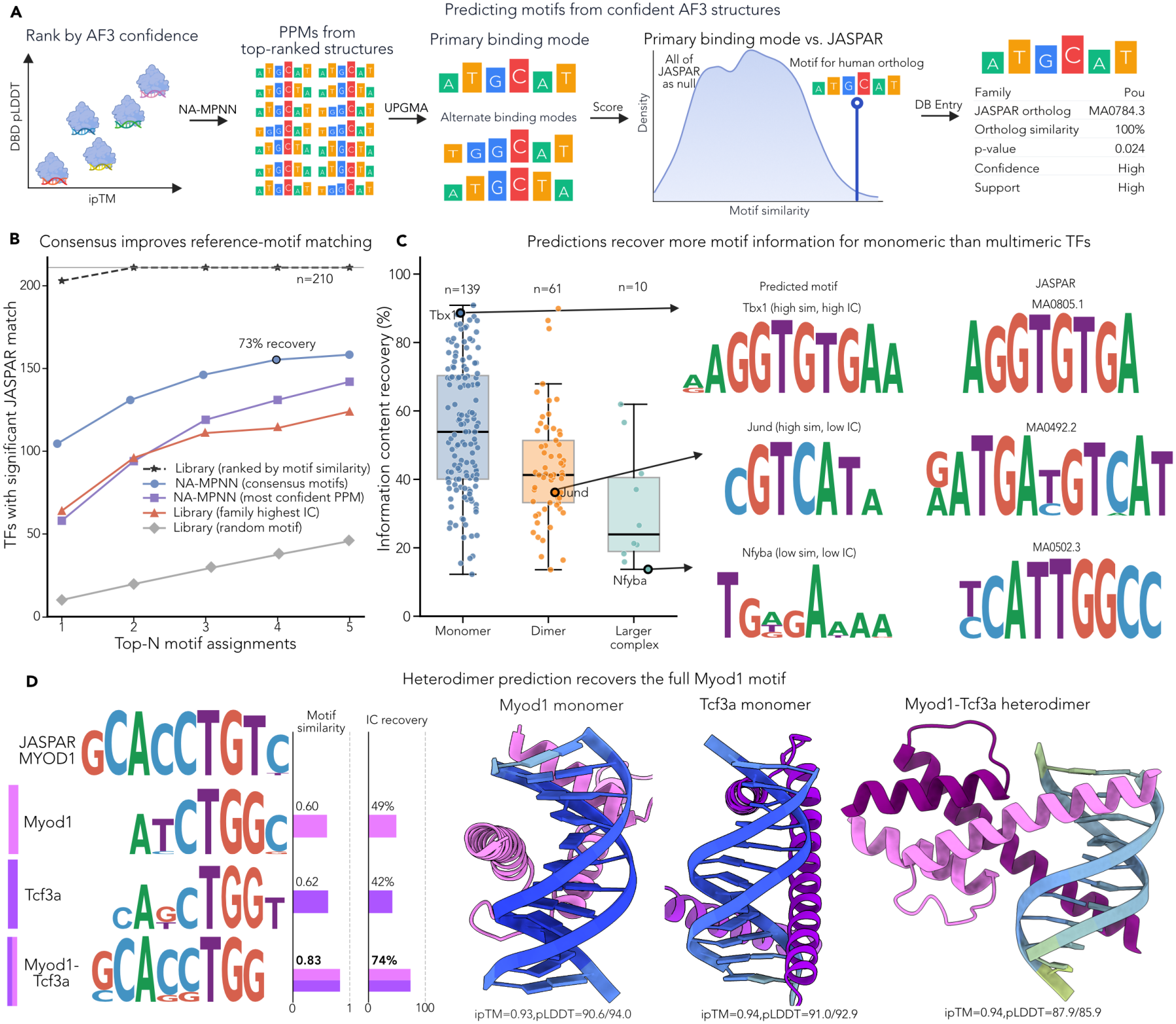
Predicted binding modes match orthologous reference motifs for two-thirds of positive controls. **(A)**Motifs are read from confident AlphaFold3 (AF3) structures and scored against the human ortholog’s reference motif. Complexes are ranked by AF3 confidence (*S*_conf_), NA-MPNN infers a position probability matrix (PPM) from each highly confident structure, and UPGMA clusters these into a primary and up to three alternate binding modes. Each mode is scored against the orthologous JASPAR motif by motif similarity, as an empirical *p*-value with all of JASPAR as null. Each database entry reports the DNA-binding domain (DBD) family, JASPAR ortholog, ortholog DBD-similarity, *p*-value, confidence and support. **(B)**Consensus across structures improves reference-motif recovery. Positive controls (*n* = 210) recovered at *p* < 0.05 versus the number of motifs assigned per TF. At the deployed configuration, consensus binding modes match 103 TFs (49%) at the primary mode and 153 (73%) within four (in-sample; 44.0% and 66.7% under cross-validation), against 64 and 114 for the family baseline (the highest-IC library motif from the TF’s DBD family) and an expected 10 and 38 for a random library draw. Assigning individual NA-MPNN PPMs from the most confident complexes without consensus recovers 58 at the primary mode. Ranking library motifs by motif similarity to the reference, a ceiling bounded by library composition, recovers 203 at the primary mode and all 210 within two. **(C)** Information content (IC) recovery declines with binding-complex size. IC recovery of the primary mode by binding-complex class (monomer *n* = 139, dimer *n* = 61, larger complex *n* = 10; medians 54%, 41%, 24%). Predicted primary modes (left) and JASPAR references (right): Tbx1, a monomer recovered at high motif similarity and IC; Jund, a bZIP homodimer whose correct half-site scores high on motif similarity but low on IC; and Nfyba, the non-sequence-specific NF-Y subunit, whose reference describes an assembly specified by another chain. **(D)** Modeling the Myod1-Tcf3a heterodimer recovers the full MYOD1 motif. Primary modes predicted for the Myod1 and Tcf3a monomers and the Myod1–Tcf3a heterodimer, aligned to the JASPAR MYOD1 motif, with motif similarity and IC recovery (bars); monomers are scored against their own JASPAR orthologs. Representative structures (Myod1 pink, Tcf3a purple) are annotated with ipTM and DBD/DNA-pLDDT, and DNA is colored by pLDDT.

**Fig. 4.**
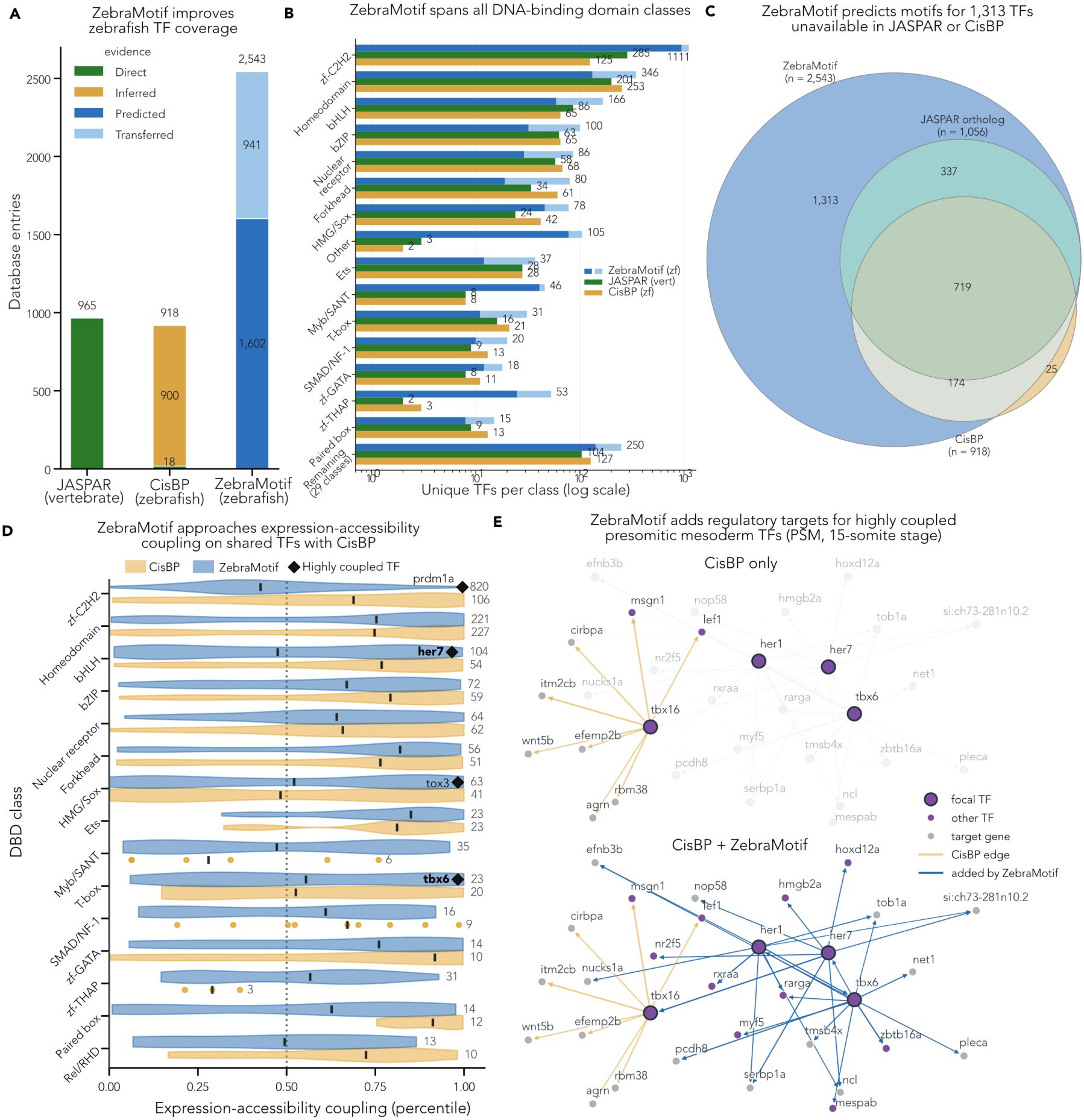
ZebraMotif, a proteome-scale structure-based motif resource for zebrafish transcription factors (TFs). **(A)** ZebraMotif covers far more zebrafish TFs than existing resources: 2,543 entries (1,602 directly predicted, 941 transferred via cluster membership) versus 918 for CisBP and 965 vertebrate PWMs for JASPAR. Bars colored by evidence type. **(B)** The gain is present in every DNA-binding domain (DBD) class and is largest for zf-C2H2. Unique TFs per class (log scale; classes reconcile AnimalTFDB4, JASPAR and CisBP family annotations) **(C)** ZebraMotif adds motifs unavailable elsewhere. Overlap of ZebraMotif (*n* = 2,543), CisBP (*n* = 918), and JASPAR orthologs (*n* = 1,056): ZebraMotif adds 1,313 TFs absent from both prior resources. **(D)** ZebraMotif motifs recover TF expression–accessibility coupling close to CisBP’s zebrafish collection on shared TFs, across DBD classes. Violins and medians show all scored TFs per resource. The statistics given in the text are paired on the 726 TFs scored in both. Per-class TF counts are listed at right (ZebraMotif above, CisBP below). In classes with too few members for a reliable density estimate, individual TFs are shown as points. Diamonds indicate four highly coupled TFs (*prdm1a, her7, tox3, tbx6*) whose ego-regulons are shown in Supp. Fig. 13B. Classes are ranked by size for the 15 largest. **(E)** ZebraMotif adds regulatory targets for highly coupled presomitic mesoderm (PSM) TFs. Networks seeded by four TFs (*tbx6, tbx16, her1, her7*; large purple nodes) from the CellOracle GRN for the PSM at the 15-somite stage, built from the CisBP-only (top) and hybrid CisBP + ZebraMotif (bottom) base GRN. Each network shows the top eight edges per seed by coefficient (*p* < 0.001), plus all significant edges among the displayed nodes. Arrows point from TF to target. Orange edges are present in the CisBP-only GRN, and blue edges appear only with ZebraMotif motifs. Small purple nodes are other TFs, and grey nodes are target genes.

The clearest failures fall into three cases (Fig. 1B). Phf21ab is a PHD-finger chromatin reader without sequence-specific DNA binding. AF3 models the THAP-domain paralogs Prkrira and Prkrirb with low confidence. Pax2b fails (AUROC 0.44), whereas its paralog Pax2a, with a near-identical paired domain but scored against a JASPAR rather than a CisBP motif, discriminates almost perfectly (0.99), which points to the reference motif being incorrect rather than the prediction.

### Discrimination is carried by interface confidence and is strongest for confidently predicted TFs

The discriminating signal is local to the protein-DNA interface (Supp. Fig. 4A). The interface also contains geometric signal, as co-folding with cognate DNA draws interface side chains closer to the nucleobases (Fig. 1C), and TFs whose interfaces are drawn closest are those that confidence discriminates best (Pearson *r* = 0.48, *p* = 0.009). Discrimination further scales with overall prediction confidence, so that the most confidently predicted TFs are those the score discriminates best (AUROC versus the mean confidence of each TF’s motif-bound structures, Pearson *r* = 0.78; Supp. Fig. 4B) and low-confidence predictions can be flagged. Because absolute discrimination varies across TFs, we asked whether confidence is useful as a ranking signal instead of acting as a hard classifier. Retaining only the top *~*4.5% of structures by confidence raises the fraction that are motif-bound from the 0.17 base rate (30 motif versus 150 non-motif structures) to a median of about 0.50, a three-fold enrichment (Fig. 1D). Confidence therefore enriches for cognate DNA even where the motif and non-motif distributions overlap.

### A proteome-scale computational screen to predict motifs for all zebrafish TFs

Having established that AF3 confidence metrics discriminate cognate from non-cognate DNA, we used this signal as a gate for a computational screen that predicts a motif for every zebrafish TF from its most confident complexes (Fig. 2A). The screen searches a DNA library of candidate motifs against a panel of TF targets, scoring each pairing by AF3 confidence. The library sequences that form the highest-confidence complexes with a target are its hits, and the target’s motif is then read from the structures of these complexes.

We built a library of zebrafish-derived binding targets by deriving candidate motifs from accessible chromatin in the developing zebrafish embryo. Zebrahub-Multiome^29^ groups over 640,000 accessible-chromatin peaks from 10–24 hours post-fertilization (hpf) embryos into 36 Leiden clusters by their accessibility across cell types and time-points, and we discovered motifs within each cluster using HOMER,^31^ capturing motifs from the distinct regulatory programs active across cell types and developmental stages (Fig. 2B; Supp. Fig. 5A). Clustering the 2,566 motifs (all *de novo* motifs plus JASPAR matches at *p* < 10^−12^) gives 312 non-redundant motifs, of which 298 came from HOMER’s *de novo* discovery and 14 were JASPAR matrices.^11^ Most *de novo* motifs resemble a known JASPAR motif without duplicating it. Their median BestGuess score (HOMER’s similarity to the closest JASPAR match) is 0.68 (Supp. Fig. 5B), and most share a core with their nearest JASPAR relative but differ in IC and flanks (Supp. Fig. 5C). The library thus supplies zebrafish-derived variants of known specificities in 312 motifs, against over 1,900 in the redundant vertebrate JASPAR collection.

Next, we retrieved zebrafish TFs from AnimalTFDB4,^30^ excluding annotated cofactors and chromatin remodelers, which left 2,543 TFs. To reduce the number of AF3 runs, we collapsed TFs expected to share DNA-binding specificity into 1,602 cluster representatives, a 37% reduction (Fig. 2C, Supp. Fig. 6A). zf-C2H2, ZBTB, and homeodomain TFs were clustered by recognition code (see Methods) because global similarity in these families is dominated by scaffold residues, and all other families by MMseqs2^32^ on their concatenated annotated domain sequences (Methods). Members of a cluster remain more similar to one another than to members of other clusters (Supp. Fig. 6B).

### A two-pass screen concentrates compute on predicting confident and geometrically resolved interfaces

In the broad first pass, every target was scored against the full motif library and its hit motifs were ranked by average ipTM per motif; the enriched second pass then sampled the most confident motifs more deeply (see Methods). The enriched pass is only *~*23% of all structures generated (482,321 of 2,117,605) yet supplies *~*41% of the most confident structures (top 5% by *S*_conf_), nearly double its share of the pool (Supp. Fig. 7B).

AF3 generates confident protein–DNA interfaces across diverse DBD families (Fig. 2D), though the confidence (mean per-TF ipTM × DBD-pLDDT) varies by family, from highest in ETS and T-box to lowest in THAP (Supp. Fig. 7A). We then tested whether interfaces resolve known contacts for C2H2 zinc finger TFs, where the recognition code provides ground truth for what residues contact DNA. Within the fingers AF3 calls as DNA-binding, the base-directed residues match the known specificity-determining positions, and the fraction of a residue’s contacts that are base-directed distinguishes base readers from backbone contacts (F1 = 0.92; Fig. 2E; Supp. Fig. 7C; Methods).

### NA-MPNN converts confident interfaces into a zebrafish motif database

A confident interface reports how well AF3 placed one DNA sequence, whereas a motif describes the range of sequences a TF prefers. We therefore took the top-ranked complexes from both passes and used NA-MPNN’s specificity mode,^24^ which predicts a position probability matrix (PPM) for the DNA from the backbone coordinates of a protein–DNA complex. Because *S*_conf_ is a within-TF rank, every TF contributed its top-ranked complexes regardless of their absolute confidence. We clustered these PPMs into a primary binding mode and up to three alternates, each reported with its single most confident structure (Methods). Where a JASPAR ortholog exists, each entry also reports its DBD similarity and the motif similarity and empirical *p*-value of the predicted mode against the ortholog’s motif (Fig. 3A).

### Predicted binding modes match the orthologous motif for two-thirds of positive controls

We bench-marked these predictions on 210 positive-control TFs whose ZFIN-curated human ortholog has a JASPAR motif and shares ≥95% DBD similarity (or recognition-code identity) with the zebrafish TF (Supp. Fig. 8; Methods). A predicted mode counts as a match when it is more similar to the ortholog’s motif than to 95% of JASPAR (empirical *p* < 0.05; Supp. Fig. 9A; Methods). Under repeated five-fold cross-validation, with hyperparameters selected on training folds and evaluated on held-out TFs, the primary mode matches the reference motif for 44.0 ± 5.0% of TFs and within the top four modes for 66.7 ± 4.9%. At the deployed configuration, whose hyperparameters were selected on all 210 controls, the corresponding in-sample rates are 49% and 73% (Fig. 3B; Supp. Fig. 9B; Methods).

To interpret these rates, we compared them with an upper bound, two baselines, and an ablation (Fig. 3B; see Methods). First, we asked whether a correct motif was available to the screen in the library, so for each TF we took the library motif with the highest motif similarity to its JASPAR reference. This best-case choice uses the answer and is an upper bound; it matches 203 of 210 controls. Second, we asked how well a TF’s motif can be predicted without the screen. Assigning a random library motif matches 5% of controls (10/210) with one motif per TF and 18% with four (38/210). Assigning the highest-IC library motif from the TF’s own DBD family, a family baseline, matches 30% (64/210) and 54% (114/210). The pipeline matches 49% and 73% (44.0% and 66.7% under 5-fold cross-validation), so it captures specificity beyond what its DBD family predicts. Third, we asked how much the consensus step contributes. Using only the NA-MPNN PPM from each TF’s single most confident complex matches 28% (58/210) at the primary mode, below the family baseline, whereas clustering PPMs from many complexes into binding modes matches 49%. Single structures are therefore unreliable readouts on their own, and the match rate depends on agreement across many. With a correct motif in the library for 203 of 210 controls, the remaining difference is set by confidence ranking and PPM readout.

### Cluster size tracks agreement w ith the reference more closely than structure confidence

The confidence of the structures in a predicted mode and the size of its cluster can both help indicate whether it is correct. Confidence correlates with accuracy both by motif similarity, a more global measure of agreement between two matrices (*ρ* = 0.43), and by IC recovery, which measures whether the signal was captured at the right positions (*ρ* = 0.38; Supp. Fig. 9C). These correlations are modest partly because only each TF’s top-ranked structures enter PPM prediction, which narrows the confidence range. That the relationship persists within the top-ranked set means confidence still indicates quality after gating. Cluster size, the number of NA-MPNN PPMs supporting a mode, is the stronger indicator (*ρ* = 0.59 against motif similarity; Supp. Fig. 9D), and it ranks modes within an entry.

### Low information content recovery often reflects reference stoichiometry rather than prediction failure

Motif similarity and IC recovery are positively correlated (Supp. Fig. 9E), but where they disagree, binding-complex stoichiometry is often the reason. An obligate dimer’s correct half-site scores high on motif similarity and low on IC recovery, as for the bZIP homodimer Jund (Fig. 3C; Supp. Fig. 9F). Dividing the TFs in the control set by binding-complex class (see Methods) shows that IC recovery falls monotonically with complex size. Monomers recover a median 54% of the reference motif’s IC, dimers 41%, and larger complexes 24% (Fig. 3C). The best-recovered monomers approach the complete reference (e.g. Tbx1, 88.6%). Nfyba is the limiting case. It is a histone-fold subunit of NF-Y without sequence specificity of its own, so its JAS-PAR reference motif is specified by NFYA, which no Nfyba prediction can recover. In these cases, the miss comes from scoring a monomeric prediction against a multimeric reference. Modeling the assembled heterodimer recovers the full-length motif in one such case (below).

A separate limitation appears within the monomers, where C2H2 zinc-finger TFs (*n* = 40) have a median IC recovery of 40%, against 54% across monomers overall. Because zinc fingers are the largest family in the resource, we return to this in the Discussion.

### Modeling a heterodimer recovers a full-length motif

IC recovery declined with complex size (Fig. 3C), consistent with a monomer engaging only part of a multimeric site. We therefore modeled the full complex for a representative case, seeding each complex with DNA from a small set of confident monomeric structures (Methods). Myod1 and Tcf3a, which bind their motif as a heterodimer, each recovered only a half-site as monomers (motif similarity 0.60 and 0.62 to their JASPAR orthologs, *p* = 0.10 and 0.12; IC recovery 49% and 42%), but jointly recovered the full MYOD1 motif when modeled together (motif similarity 0.83, *p* = 0.029; IC recovery 74%; Fig. 3D). ipTM was unchanged (0.93 and 0.94 as monomers, 0.94 as a heterodimer), while DBD-pLDDT and DNA-pLDDT decreased modestly (from 90.6–91.0 to 87.9, and from 92.9–94.0 to 85.9; Fig. 3D).

To test whether the approach extends beyond dimers, we modeled a Tp63–Tp73 tetramer on sequences built by duplicating the palindromic sites used for the two homodimers (Methods). The predicted PWM spanned the full-length TP63 motif and matched it by motif similarity (0.88, *p* = 0.006), but recovered only 22% of its IC, less than the Tp63 and Tp73 homodimers recovered from their half-sites (24% and 33%; Supp. Fig. 10A). Confidence fell sharply from the homodimers to the tetramer (ipTM 0.86–0.87 to 0.41), with most of the decrease in DNA placement and less in protein fold (DNA-pLDDT 60–66 to 30, against DBD-pLDDT 90 to 82; Supp. Fig. 10B). AF3 struggled to position the DNA against four protein chains at once, and the approach does not yet extend reliably to assemblies larger than dimers.

### ZebraMotif expands zebrafish motif coverage across every DBD clas

ZebraMotif (zebramotif.apps.czbiohub.org) contains entries for 2,543 zebrafish TFs, 1,602 directly predicted and 941 transferred from their cluster representative (Fig. 4A). For two zf-C2H2 and two homeobox clusters, motifs predicted independently for each member match the representative’s prediction, with a single exception (Egr2a; Supp. Fig. 11A,B; Methods), supporting transfer of the representative’s motif to the 941 non-representative TFs. The total coverage is 2.8× the coverage of CisBP (918 TFs), the largest existing zebrafish motif collection, in which 18 entries come from direct experiments and the other 900 are assigned through homology transfer from other species. JASPAR holds 965 unique vertebrate PWMs with direct experimental evidence but assigns none to zebrafish (Fig. 4A). The gain is present in every DBD class and is largest for C2H2 zinc finger TFs, where ZebraMotif covers 1,111 TFs, nearly nine-fold CisBP’s 125 (Fig. 4B). CisBP (*n* = 918) and JASPAR (*n* = 1,056, counted by ortholog mapping from ZFIN where one PWM can account for multiple paralogs) overlap heavily with each other, whereas ZebraMotif adds 1,313 TFs absent from both databases (Fig. 4C). ZebraMotif lacks 25 TFs that CisBP covers. Most of them (*n* = 17) are chromatin regulators, such as CXXC-domain CpG readers, and are not labeled as sequence-specific TFs in AnimalTFDB. The remaining 8 include general transcription factors (Tbp, Tbpl2) and a few canonical sequence-specific TFs absent from AnimalTFDB4 (e.g. Pparg, Esrrd).

Each entry can be browsed in a web interface that displays the primary and alternate binding modes, their cluster support and similarity to JASPAR orthologs, and the highest-ranked AF3 structure with its confidence scores (Supp. Fig. 14; Supp. Video 1).

### ZebraMotif approaches CisBP’s expression– accessibility coupling on shared TFs

To test whether ZebraMotif motifs are functional in zebrafish chromatin, we measured the coupling between each TF’s expression and the accessibility of its predicted binding sites. A TF’s expression and the accessibility of its sites are coupled across cells, whether the TF opens chromatin itself or binds and stabilizes sites already primed,^33^ and this coupling is routinely used to identify the TFs driving cell states.^34, 35^ Because coupling is recovered only if the motif points to the sites the TF occupies, it offers a reference-free, class-level check on motif quality. For each TF, we scored how well its expression tracks accessibility at its predicted sites across the Zebrahub-Multiome atlas, ranked against every other TF’s expression (Supp. Fig. 12A; Methods). For example, *otx2b* expression correlates only modestly with accessibility at its predicted sites (Spearman *ρ* = 0.22), but this correlation is higher than that of the same sites with 99% of other TFs (Supp. Fig. 12B). Coupling depends on confounding variables such as motif IC and paralogs, so we used it comparatively, benchmarking ZebraMotif against CisBP on the same TFs and within each DBD class (Fig. 4D). On the 726 TFs scored in both resources, ZebraMotif motifs couple slightly less strongly than CisBP’s (median paired difference 2.6 percentile points; Wilcoxon signed-rank *p* = 1.2 × 10^−8^), and CisBP is significantly higher than ZebraMotif in three of the 15 largest DBD classes: paired box (by 15.9 points), zf-C2H2 (10.9) and bZIP (5.1; *q* < 0.05, Benjamini–Hochberg).

zf-C2H2 TFs are the hardest class for the screen, likely because their long finger arrays can engage DNA in many registers (Discussion). Consistent with this, the gap extends beyond shared TFs. Across all 820 zf-C2H2 TFs with a ZebraMotif motif, most of which have no motif in CisBP, median coupling (43rd percentile) falls just below the TF-swap null (50th) (Fig. 4D). Together with their lower reference-motif recovery (Fig. 3C), this suggests treating ZebraMotif’s zf-C2H2 entries as candidates to prioritize by their confidence and cluster support, rather than as confident assignments.

### ZebraMotif adds candidate regulatory edges consistent w ith know n segmentation biology

Gene regulatory network (GRN) inference is a primary use of a motif database. Motifs link each TF to the accessible regions it may bind and, through them, to nearby genes. We inferred cell-type- and timepoint-specific GRNs with CellOracle^36^ from CisBP-only and hybrid (CisBP + ZebraMotif) base GRNs, changing nothing else (Supp. Fig. 13A; Methods). In the 15-somite presomitic mesoderm (PSM), we seeded the network with four TFs central to segmentation, *tbx6, tbx16, her1* and *her7* (Fig. 4E), including the highly coupled *tbx6* and *her7* (Fig. 4D). Only *tbx16* has a CisBP motif, so in the CisBP-only network it is the only seed with targets. With ZebraMotif, all four regulate targets and the seeds become linked, including *tbx6* →*her1, tbx6* →*her7* and *her7*→*tbx16*.

The added edges agree with prior genetic and cis-regulatory data. The top targets of *tbx6* include *her1, her7, pcdh8* and *mespab*, all with activating coefficients (Supp. Fig. 13B). Tbx6 (formerly Tbx24, the *fused somites* gene product) binds a T-box site in a *her1* upstream element in vitro, and this site is required for reporter expression in the anterior PSM;^37^ Tbx6 also activates the *her1* upstream region in a cell-based reporter assay.^38^ In *tbx6* mutants, the most anterior *her7* stripe is lost while posterior oscillation persists,^38, 39^ *pcdh8* is lost from the anterior PSM,^40^ and *mespab* expression is absent.^41^ Edges from two other highly coupled TFs, *tox3* in the PSM and *prdm1a* in the epidermis, have no prior characterization and are reported as candidates (Supp. Fig. 13B). Because each network is inferred within a single cell state, regression cannot separate direct regulation from co-expression, so we treat the added edges as consistency checks on the predicted motifs, not validated interactions.

## Discussion

We present a structure-based screen that infers DNA-binding specificity for every TF in a proteome by predicting TF–DNA complexes with sequences sampled from candidate motifs drawn from the organism’s own accessible chromatin. In zebrafish, a two-pass design concentrated compute on high-confidence complexes, and NA-MPNN inferred motifs from their DNA-bound structures. The resulting TF-motif database, ZebraMotif, covers 2,543 annotated TFs.

The screen relies on two signals: AF3 confidence at the protein–DNA interface, which enriches for cognate DNA, and the second is agreement among many confident complexes. AF3 assigned higher confidence to cognate than to non-cognate DNA for most TFs across 20 DBD families, with the clearest failures in a protein without a canonical sequence-specific DBD, in TFs that AF3 models with low confidence, and in one paralog whose reference motif likely differs from its true specificity. Confidence can therefore act as a gate, and a two-pass design that resamples each TF’s most confident motifs, loosely analogous to successive rounds of SELEX enrichment, makes a proteome-scale search computationally tractable.

Confident structures are not yet motifs. Contact geometry tracks discrimination only partially, and each complex captures a single sequence that AF3 placed confidently, whereas a motif describes the range of sequences a TF prefers. NA-MPNN infers a motif from each complex without family-specific recognition rules, and recovery of the reference motif depends on consensus across many complexes; the number of NA-MPNN PPMs supporting a mode is also a better indicator of its reliability than structure confidence.

Our approach requires no binding assays and little prior data. Here, the candidates came from zebrafish accessible chromatin, but where accessibility data are unavailable, existing collections such as JASPAR could supply the candidate library. Readout methods such as DeepPBS,^23^ and NA-MPNN as used here, infer specificity from a protein–DNA complex, but building that complex requires knowing which DNA the TF binds. Our screen searches for that DNA, which lets it reach TFs that current approaches miss. Consortium-scale assays now cover most human TFs,^8^ and homology transfer extends these measurements to species with similar DBDs.^10, 12^ Neither covers TFs whose domains have diverged too far for transfer, or species where experiments at this scale are unlikely. In zebrafish, ZebraMotif adds motifs for 1,313 TFs absent from both CisBP and JASPAR, and on TFs shared with CisBP, its motifs show expression–accessibility coupling close to CisBP’s. Each motif also comes with a predicted structure that can point to residues for mutagenesis or protein design, and with confidence and cluster support that indicate how reliable it is.

Our approach has four main limitations. First, the benchmark includes only TFs with a close human ortholog; performance on TFs with diverged DBDs, the cases the screen is meant for, is untested and requires experimental validation. The bench-mark also cannot separate inference from recall, as both AF3 and NA-MPNN were trained on experimental protein–DNA complexes, which are enriched for the well-studied families that make up our positive controls, so match rates for TFs without structural precedent may be lower. Second, motif predictions were made for monomeric TFs, whereas many TFs bind as multimeric complexes that add or change specificity around a core, and IC recovery in the benchmark declines with complex size. Seeding a Myod1–Tcf3a heterodimer with DNA from confident monomeric structures recovered the full-length human MYOD1 motif, but this was a proof of concept, and predicting multimeric structures systematically is a combinatorial problem that warrants its own screen design. Third, C2H2 zinc-finger TFs recover less of the reference IC than other monomers and show weaker expression– accessibility coupling with our predicted motifs than with CisBP’s, possibly because DNA is placed in the wrong register or the screened sequences are too short to accommodate every zinc finger that may engage *in vivo*. Fourth, every annotated TF receives an entry, although not all may bind DNA sequence-specifically, and entries transferred from a cluster representative were not predicted directly; each entry reports confidence and cluster support to indicate reliability.

HT-SELEX is underway on several TFs without characterized motifs and will test the method where no ortholog evidence exists. For C2H2 zinc fingers, recognition-code predictors such as DeepZF^42^ could help identify the engaged fingers and seed predictions with DNA that is length-appropriate for the register. The screen reported here used *~*107,000 GPU-hours; faster structure prediction models like ESMFold2^22^ and DBD-only inputs would lower the cost per prediction and allow denser sampling of the search space. The pipeline requires only a proteome and a set of candidate motifs, neither of which depends on prior binding data for the TFs being predicted, so it can be applied to other species with an annotated genome without waiting on experimental throughput.

## Supporting information

Supplementary Video 1. Walkthrough of the ZebraMotif website

## Acknowledgements

We thank the Computational Biology platform and the members of the Royer and DeRisi labs for helpful discussion and support. We thank the Scientific Computing team for their support in running our large-scale prediction pipeline. Funding for this work was provided by Biohub. We thank the Biohub donors, Priscilla Chan and Mark Zuckerberg, for their generous support.

## Author contributions

J.L.D. and L.A.R. conceived the project. B.I., Y.J.K., and L.A.R. developed the idea into the computational screen. B.I. built the screening and motif-prediction pipeline, ran the predictions, performed the analyses, and built the ZebraMotif website. W.L., N.H., M.P., and R.A. deployed AlphaFold3 on the HPC cluster and helped with GPU scheduling for the large-scale AF3 predictions. N.B., L.B., and J.P. designed and are carrying out the HT-SELEX experiments. Y.J.K. and L.A.R. supervised the project. B.I., Y.J.K., and L.A.R. wrote the manuscript with input from all authors.

## Data availability

ZebraMotif version 0.1 and a walkthrough video of the website (Supp. Video 1) are available at Zenodo (doi.org/10.5281/zenodo. 23380219). The website is accessible at zebramotif. apps.czbiohub.org. Code availability Code for the screen, PPM prediction and benchmarking is available at github.com/ czbiohub-sf/tf-motif-screen. Code for the website is available at github.com/czbiohub-sf/zebramotifexplorer.

## Methods

### Transcription factor selection for motif–non-motif experiments

We assembled a benchmark set of 29 zebrafish TF binding profiles from CisBP (v 3.10)^10^ and JASPAR (2024).^11^ Of these, 16 TFs from CisBP have direct experimental evidence from protein binding microarray (PBM) assays, and another 13 were selected from JASPAR by identifying zebrafish TFs with ≥95% DNA-binding domain (DBD) sequence similarity to a human ortholog (see Determining orthology between zebrafish and human transcription factors), expanding coverage across large DBD families (Supp. Fig. 1A). For each family, we characterized the AF3 database-search outputs by counting the sequences in each TF’s paired multiple sequence alignment (MSA), measuring DBD sequence identity to the returned protein templates, and recording whether the returned PDB entries included DNA. These quantities were summarized within the 20 DBD groups shown in Supp. Fig. 1A. Selected DBD superpositions and a comparison with a full-protein superposition are shown in Supp. Fig. 1B,C.

### Motif–non-motif experiments

To test whether AF3 distinguishes cognate from non-cognate DNA, we paired each TF with DNA sequences sampled from its position weight matrix (PWM) (positives), as well as length- and GC-matched genomic non-motif sequences (negatives) (Supp. Fig. 2A). Non-motif sequences were drawn as k-mers of the same width as the motif from accessible-chromatin peaks. Each candidate had to score below the PWM’s *p* < 10^−3^ log-odds threshold and contain none of the motif’s high information content (IC) core seeds on either strand. This produced negatives with realistic genomic base composition that controlled for sequence-specific recognition, GC content, and length. Each protein-DNA pair was predicted with AF3 as a single TF chain and its DNA sequence. We compared the confidence distributions of motif and control complexes within each TF.

To determine the number of sequences to sample per PWM, we sampled varying numbers of sequences from each PWM (*P*), reconstructed a PWM (*P*^′^) by position-frequency counting, and measured reconstruction accuracy as the Jensen-Shannon divergence (JSD) between *P* and *P*^′^ (Supp. Fig. 2B). JSD decreased as more sequences were sampled (Supp. Fig. 2C) and saturated near *n* = 30 for most PWMs (Supp. Fig. 2D). We therefore sampled 30 sequences per PWM as the resolution for all discrimination analyses. We sampled 150 non-motif sequences per TF, more than the 30 motif sequences, to define the null against which motif structures are ranked. Under this design a random ranking has an average precision of 0.17, so values above this base rate indicate motif-bound complexes are being enriched.

### Systematic determination of DNA-binding domains

We annotated TF sequences with Inter-ProScan 5.73-104.0,^43^ using Pfam 37.2 together with PROSITE profiles 2023_05. Overlapping annotation calls were merged (Supp. Fig. 3A). To identify when a domain is DNA-binding, we took the AF3 predicted structures and counted, at each protein residue, the DNA nucleotides in chains B and C with predicted aligned error (PAE) < 5 Å when the prediction is aligned on the nucleotide, using confidence in relative positioning as a proxy for the DNA-binding region. We retained residues whose counts were at least the mean among residues with nonzero counts (Supp. Fig. 3B,C). For single-domain assignments, we selected the annotated domain with the highest mean frequency of retained positions across the analyzed predictions, averaging over the domain’s residues. TFs in the zf-C2H2 and ZBTB families could retain multiple domains because they can have more than one DNA-binding C2H2 zinc finger.

### Evaluating discrimination from AF3 confidence values

We evaluated predictions using three AF3 confidence metrics. The interface predicted template modeling (ipTM) is a score in [0, 1] reflecting AF3’s confidence in the geometry of the interface between chains. We used the protein–DNA ipTM, the mean of AF3’s chain-pair ipTM between the protein chain and each DNA strand (for multi-chain complexes, the protein chain with the highest value). The predicted local distance difference test (pLDDT) is a per-atom score in [0, 100] reflecting confidence in local structural accuracy. We computed two summaries, DBD-pLDDT (the mean pLDDT over the identified DBD after adjusting its boundaries to the confidently folded region, extending outward while flanking residues have pLDDT > 70 and trimming inward while edge residues have pLDDT < 70; predictions with no DBD residue above 70 were excluded, and TFs with no assigned DBD were given a DBD-pLDDT of zero) and DNA-pLDDT (the mean pLDDT across all nucleotides in the DNA chains). PAE is a pairwise score, in Ångstroms, reflecting AF3’s confidence in the relative positioning of two residues or nucleotides. Interface PAE is AF3’s minimum PAE between the protein chain and each DNA strand (chain_pair_pae_min), averaged over the two strands. Because no metric or pair discriminated substantially better than another (Supp. Fig. 4C), we combined ipTM with DBD-pLDDT, which discriminates TFs that interface metrics miss (Supp. Fig. 4D) and whose product also predicts per-TF discrimination (Supp. Fig. 4B), so that unreliable TFs can be flagged. Each complex *i* in a pool of *N* complexes is scored as

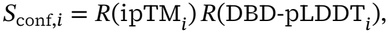

where *R*(*x*_*i*_) is the ascending rank of *x*_*i*_ divided by *N*, with ties given average ranks. For example, in the motif–non-motif benchmark the pool was all retained predictions for a TF (up to 180). Where a raw score is shown (Fig. 1B; Supp. Figs. 4B, 7A and 9C) we use ipTM × DBD-pLDDT/100.

To assess whether AF3 distinguishes motif from non-motif sequences for each TF, we scored the separation between each TF’s motif and non-motif structure distributions with two ranking metrics: the area under the receiver operating characteristic curve (AUROC) and the average precision (AP), the area under the precision–recall curve with motif as the positive class, whose random baseline equals the motif fraction (30 motif versus 150 non-motif sequences per TF, i.e. 0.167). Both were computed per TF and macro-averaged across the 29 TFs, and per-TF significance was assessed with one-sided Mann–Whitney U tests for higher motif scores (*p* < 0.05). To choose a ranking metric, we compared per-TF discrimination across metrics that report confidence relative to the interface of the TF-DNA complex (Supp. Fig. 4C), including ipTM, DBD-pLDDT, DNA-pLDDT, interface PAE, and pTM as a background, and their pairwise combinations. We characterized how the metrics relate by correlating each metric’s per-TF AUROC with that of ipTM (Supp. Fig. 4D), and confirmed that confidence is predictive of discrimination, a prerequisite for accepting or rejecting predictions on confidence, by correlating each metric with per-TF discrimination.

### Extracting candidate DNA sequences from Zebrahub-Multiome snATAC-seq

To generate candidate DNA sequences for our systematic DNA-binding motif screen, we extracted over-represented sequence motifs from a dataset of over 640,000 peaks from accessible chromatin in 10–24 hours post-fertilization (hpf) zebrafish embryos.^29^ We processed peaks per cluster, using Zebrahub-Multiome’s Leiden clustering of peaks by accessibility profiles across time points and cell types, with each cluster representing a potential regulatory program. We ran HOMER^31^ on DNA sequences from each cluster with default parameters, queried against the JASPAR 2024 CORE vertebrates (redundant) database, and mapped to genome assembly GRCz11 (Supp. Fig. 5A). JASPAR-matched motifs were retained at *p* < 10^−12^ to ensure confident assignments; all *de novo* motifs were retained regardless of significance to maximize sequence diversity for the downstream screen. To reduce redundancy, all resulting motifs were clustered using HOMER at its default correlation threshold of 0.6, reducing 2,566 motifs to 312 non-redundant candidates.

To characterize how closely the *de novo* motifs correspond to known motifs, we plotted the BestGuess score of each *de novo* motif (*n* = 298; Supp. Fig. 5B) and a representative *de novo* motif at the median score alongside its BestGuess match (Supp. Fig. 5C). BestGuess scores are taken as reported by HOMER and are used only to characterize and cluster the candidate library; all other motif comparisons use motif similarity (Motif similarity).

### Recognition codes

Two DBD types read DNA through a small number of known specificity-determining residues, and we used those residues directly wherever specificity had to be compared instead of relying on global sequence similarity. For C2H2 zinc fingers, all zinc finger repeats were identified using a regular expression anchored on the conserved C2H2 spine (C.{2,4}C.{3}[FYL].{5}[LIVMFY].{2}H.{3,5}H), where the hydrophobic-core positions are matched as residue classes rather than the canonical F/Y and L/I, and each finger was reduced to the four DNA-contacting positions p−1, p2, p3, and p6 relative to the recognition helix.^17^ For homeodomains, sequences were extracted from full-length proteins using InterProScan boundary annotations (IPR001356/PF00046), aligned to the Pfam PF00046 HMM^44^ using pyhmmer’s hmmalign,^45^ and reduced to a five-character code at the canonical DNA-contacting positions 47, 50, 51, 52, and 54.^16^ These codes are the basis of clustering (Clustering transcription factors), of ortholog similarity (Determining orthology between zebrafish and human transcription factors), and of the ground truth used to calibrate base-reader calling (Protein-DNA contact extraction and base-reader calibration). Positions entered the code because the recognition literature assigns them a base-reading role. A coded position is not guaranteed to fall within contact distance of DNA in every structure, and homeodomain position 52 systematically does not (Protein-DNA contact extraction and base-reader calibration).

### Clustering transcription factors

We took zebrafish TF sequences from AnimalTFDB4.0,^30^ excluding entries labeled as chromatin remodelers or cofactors, which left 2,543 sequences in 72 families. To reduce redundant AF3 runs while preserving DNA-binding diversity, we clustered TFs within each family. Zinc-finger C2H2, homeodomain, and ZBTB TFs were clustered by recognition code (Supp. Fig. 6A; Recognition codes), because global similarity in these families is dominated by scaffold residues. All other families were clustered with MMseqs2 easy-cluster^32^ (default settings) on each protein’s annotated domain sequence, so that clustering reflects domain content rather than linkers or disordered regions. Cluster representatives were folded as fulllength proteins.

For zf-C2H2 and ZBTB TFs, proteins were partitioned by finger count and pairwise recognition-code distances were computed: positions p−1 and p3 required exact identity (a mismatch at either voided the finger and set the protein-pair distance to infinity), while p2 and p6 contributed one unit per mismatch (maximum 2 per finger). Proteins within the same ZF-count group were clustered by average-linkage hierarchical clustering with a threshold of one mismatch per ZF. For each cluster, the representative was the medoid minimizing summed recognition-code distance to all other members. Ties were broken by the number of InterProScan-annotated C2H2 domains, then alpha-betically by Ensembl gene identifier.

Homeodomain TFs were clustered by identity across their five-position recognition code (Recognition codes). Proteins were grouped by secondary domain context (POU, PAX, LIM-HD, CUT-HD, SIX, HNF1, ADNP, or SOLO) to account for domain-context-dependent binding specificity. Within each class, proteins were clustered greedily: two sequences were co-clustered if all five canonical positions were gap-free and identical. The cluster representative was chosen as the medoid: the member with the highest mean pairwise percent identity across all HMM match-state columns.

To assess within-cluster redundancy, sequences within each cluster (clusters with more than one sequence) were aligned using MAFFT (–auto mode)^46^ and all pairwise similarities were computed from the resulting multiple sequence alignment (Supp. Fig. 6B). To quantify inter-cluster similarity, sequences from each pair of clusters were pooled and co-aligned using MAFFT, and pairwise similarities were computed across cluster boundaries only. For each pair of aligned sequences, similarity is defined as the fraction of aligned positions receiving a positive BLOSUM62 score, excluding columns where both sequences contain a gap; gaponly positions in one sequence penalize the denominator but not the numerator. Similarity is reported as a percentage.

### Large-scale AlphaFold3 predictions

We predicted every target–motif pairing between the 312-motif candidate library (Extracting candidate DNA sequences from Zebrahub-Multiome snATAC-seq) and the 1,602 cluster representatives (Clustering transcription factors) with AF3^19^ in two passes. Each DNA input was a duplex of the motif’s width, sampled from the PWM. In the broad first pass, each of the 1,602 targets was predicted against all 312 motifs with three sequences per motif (*~*1,000 predictions per target; 1,635,284 structures), then ranked per target by mean ipTM (ties broken by mean DBD-pLDDT) to retain the top 50 motifs; *S*_conf_ was used for all subsequent ranking. Three sequences per motif were sufficient because this pass ranked motifs coarsely, whereas the benchmark used 30 to reconstruct the source PWM. In the enriched second pass, each retained motif was sampled with at most ten additional sequences (482,321 structures). For every prediction we recorded pTM, ipTM, DBD-pLDDT, DNA-pLDDT, and PAE. No DNA sequence was included in more than one job, so not all motifs generated a total of 10 sequences in the second pass. Early in the screen, some TFs received more than three sequences per motif in the first pass, and these predictions were retained.

### Protein-DNA contact extraction and base-reader calibration

For each predicted complex, a residue was called a DNA contact when any side-chain heavy atom lay within 5 Å of a DNA heavy atom (main-chain N, CA, C, O excluded), and the contact was classified as backbone if the nearest DNA atom belonged to the sugar–phosphate set (P, OP1/OP2, O5’–C1’) and base otherwise. Pooling each TF’s structures gave, per residue, a contact frequency, the base-directed fraction of its contacts, and a median side-chain pLDDT. A residue was called a base reader when all three exceeded minimum thresholds.

We calibrated the thresholds on 756 zf-C2H2 TFs (*~*796,000 structures), whose base-reading positions are known from the recognition code (Recognition codes). The calibration set contained 8,072 coded positions across 2,018 DNA-binding fingers. Scoring was restricted to residues within those fingers, so negatives are the hard cases (the Cys/His spine and non-reading positions). Over a 100-point grid (contact frequency 0.1–0.5, base fraction 0.2–0.6, pLDDT 50–80), F1 peaked at 0.1/0.3/50 (precision 0.968, recall 0.877, F1 0.921, MCC 0.88), and only the base-fraction threshold shifted the precision–recall balance (Supp. Fig. 7C). Transferred unchanged to homeodomains (64 TFs, *~*65,000 structures), with position 52 removed from the positives (it points away from the groove) and the N-terminal arm excluded (its minor-groove contacts are not in the code), the thresholds gave F1 0.787 (precision 0.761, recall 0.815), within 0.005 of the homeodomain’s own optimum, so they are not zinc-finger-specific. Base-reading residues called with these thresholds are reported for every ZebraMotif entry.

### Determining orthology between zebrafish and human transcription factors

To assign ground-truth DNA-binding motifs to zebrafish TFs, each TF was matched to its human ortholog using ZFIN’s curated orthology ^47^ and the similarity of that pairing scored with a family-appropriate metric. When ZFIN assigns several human orthologs to a zebrafish TF, an ortholog with an assigned JASPAR motif was preferred with ties broken on similarity. For most TF families, full-length zebrafish protein sequences were searched against a database of all human RefSeq proteins (NCBI RefSeq; downloaded April 2026)^48^ using BLASTP^49^ with an E-value threshold of 1 × 10^−5^ (word size 3, BLOSUM62 substitution matrix, all other parameters at default). The alignment to the ZFIN ortholog was parsed to compute percent sequence similarity restricted to the annotated DBD, counting positions that are identical or score positively under BLOSUM62.

For three families, zf-C2H2, ZBTB, and homeodomain, where specificity is largely encoded by a small number of DNA-contacting residues, similarity was computed from the recognition code (Recognition codes) instead. For zf-C2H2 and ZBTB TFs, only the fingers annotated as DNA-binding from AF3 protein-DNA PAE (Systematic determination of DNA-binding domains) were scored. This set of fingers was slid along the ortholog’s full array at every register, preserving the spacing between non-adjacent binding fingers, and the best-scoring register was retained. Each finger contributes the four DNA-contacting positions (p−1, p2, p3, p6), and the similarity is matches/(4 × *n*_fingers_). A finger differing at p−1 or p3 scores 0 of 4, since a substitution at either changes the base being read, as does a finger with no orthologous match at that register. For homeodomain TFs, similarity is the percent of the five coded positions matching the ortholog’s code (5/5 is 100%, 4/5 is 80%). Per-family mean similarities are shown in Supp. Fig. 8A.

### Motif similarity

All comparisons between predicted PPMs, and between predicted PPMs and reference motifs, used motif similarity, a HOMER-style correlation^31^ that we computed ourselves so that the alignment behind each score was available for building consensus PPMs. Motif similarity is the Pearson correlation between two frequency matrices centered on the uniform background (each entry minus 0.25) and flattened over their overlapping positions. Each overlapping position contributes the dot product of the two centered rows, and the denominator is the geometric mean of the summed squared deviations over the same window. The score is maximized over all offsets with ≥6 overlapping positions and both DNA orientations, and the maximizing offset and orientation define the alignment used to build consensus PPMs (Predicting motifs for all zebrafish TFs). Because motif similarity and the BestGuess score (HOMER’s similarity to the top-scoring match) are computed by different implementations, values are not compared across the two.

### Predicting motifs for all zebrafish TFs

For each TF, predicted protein–DNA structures were ranked by decreasing *S*_conf_ (Supp. Fig. 9A), and the highest-confidence structures were submitted to NA-MPNN^24^ in specificity mode, which predicts one position probability matrix (PPM) per structure. We clustered these PPMs to summarize each TF’s predicted DNA binding and ranked the clusters by size to define the *primary binding mode* and up to three *alternate binding modes*.

Individual PPMs were clustered per TF by UP-GMA on pairwise motif similarity (Motif similarity). Inter-cluster similarities were updated with the Lance–Williams formula, and merging continued while the highest similarity between any two clusters exceeded the similarity threshold. Clustering used the top 47 PPMs (by *S*_conf_) and a similarity threshold of 0.73. The cluster medoid was the member with the highest summed similarity to all other members. A consensus PPM was formed by aligning each member to the medoid’s coordinate frame, using the offset and orientation that maximize motif similarity, and averaging probabilities position-by-position, with non-overlapping positions retaining the medoid’s values.

### Positive-control benchmark and validation of predicted PPMs

Because no proteome-scale experimental motif set exists for zebrafish, we validated predicted PPMs against zebrafish TFs whose predicted specificity can be checked against a motif transferred from a human ortholog. Using the family-appropriate ortholog similarity defined above (Determining orthology between zebrafish and human transcription factors), we accepted an ortholog’s JASPAR_11_ motif as ground truth only when DBD conservation reached ≥95% (Supp. Fig. 8B). Similarity for recognition code families allows at most one recognition-code mismatch per five zinc fingers, and requires an exact code match for homeodomains and short arrays, where a single specificity-residue substitution is more likely to reassign the base being read. TFs lacking a curated ortholog, or whose best ortholog fell below this cutoff, were excluded. The benchmark was frozen at 210 TFs before hyperparameter selection.

To validate a prediction, we compared each TF’s consensus PPM against its assigned JASPAR ortholog motif. Supplementary Fig. 9A illustrates the workflow on Sox2. Because raw similarity scores are not comparable across motifs of differing length and IC, we computed an empirical *p*-value per PPM as the fraction of all JASPAR motifs whose motif similarity to the predicted PPM equals or exceeds that of the matched ortholog. This uses the full JASPAR database as a per-PPM null distribution, asking whether the predicted PPM resembles the correct motif more than it resembles JASPAR at large, with a prediction scored as a match at *p* < 0.05. We evaluated the match rate across these controls (Fig. 3B).

Because motif similarity scores only the aligned positions, and can be suppressed by low-information flanking positions, we also computed an IC-weighted recovery fraction. At each aligned position, the overlap between predicted and reference base distributions (the sum of per-base minima) was rescaled so that a uniform prediction scores 0 and a perfect one scores 1, weighted by the reference IC at that position, and summed; the denominator is the total IC of the JASPAR reference. This quantifies what fraction of the reference motif’s IC is recovered. Reference positions outside the alignment contribute zero, so a prediction that covers one half-site of a dimeric reference recovers at most about half its IC (Supp. Fig. 9E, computed for primary binding modes). Examining TFs distributed across the space of motif similarity and IC recovery illustrates characteristic failure modes, including cases where global similarity is low but high-IC core positions are partially recovered (Supp. Fig. 9F).

### Sweeping parameters across PPM prediction space

NA-MPNN sampling and consensus parameters were selected by grid sweeps over the 210 positive controls, maximizing the number of TFs with a significant JASPAR match within the top four modes, with ties broken by higher mean IC recovery and then by smaller parameter values. Across NA-MPNN passes per structure ({1, 5, 10, 20, 30}) and temperatures ({0.1, … 0.5}), each scored by its best result over a coarse consensus grid (pool size 10–30), match counts varied little (141–157 TFs), whereas IC recovery declined with temperature (Supp. Fig. 9B, left). We therefore fixed the temperature at 0.1 and used 30 passes, the largest value, to reduce sampling noise. We then swept PPM-pool size (10–100) and similarity threshold (0.40–0.90 in steps of 0.01) (Supp. Fig. 9B, right). The maximum (157 TFs; IC recovery 0.526) and the deployed configuration (153 TFs; 0.540; *S*_conf_, 47 PPMs, threshold 0.73) differ by less than one cross-validation standard deviation, so we deployed the latter for its higher IC recovery.

To estimate out-of-sample accuracy, we repeated this second sweep under five-fold cross-validation over TFs (five repeats, seeds 0–4), selecting on the training folds by the rule above and scoring the held-out TFs. Primary-mode and top-four match rates were cross-validated separately, each selecting on its own match rate, and we report the mean ± s.d. over the 25 held-out evaluations. These rates estimate the out-of-sample performance on the strict selection rule, conditional on the NA-MPNN setting fixed in the first sweep. On all 210 controls that rule selects the 157-TF configuration rather than the deployed one, so the deployed configuration’s rates (49% and 73%) are in-sample; given the four-TF difference, the cross-validated rates remain a close estimate for it.

### Match rate baselines

We compared the pipeline’s match rate with two reference-blind baselines, an ablation, and a library ceiling. Each arm assigns *s* motifs per TF (*s* = 1–5; Fig. 3B), and a TF counts as matched if any of its *s* motifs reaches a JASPAR-null empirical *p* < 0.05 under motif similarity (Positive-control benchmark and validation of predicted PPMs). We report *s* = 1 and *s* = 4 to match the primary and top-four modes. Because every arm is scored at the same depth on the same 210 TFs, the multiplicity at *s* = 4 affects all arms equally. The random baseline is the analytic expected match rate for *s* motifs drawn uniformly without replacement from the 312-motif library, averaged across TFs, so reported counts are these expectations rounded to integers, not the outcome of a single draw. The family baseline assigns the *s* highest-IC library motifs from the TF’s DBD family, using IC as a reference-blind proxy for motif quality. Library motifs were assigned to AnimalTFDB4_30_ families through their HOMER BestGuess annotation, by gene symbol where available and by JAS-PAR identifier otherwise. The 12 TFs in families with no assigned motif were scored as failures. The consensus ablation assigns the individual NA-MPNN PPMs from the *s* highest-*S*_conf_ complexes, without clustering or consensus averaging. The ceiling is the best match rate any selection rule could achieve over this library: it assigns the *s* library motifs most similar to each TF’s JASPAR reference. Because it uses the reference, it is an oracle. It is also close to a perfect match rate, since the best of 312 diverse matrices clears *p* < 0.05 for most well-defined references.

### Binding-complex class assignment

The IC a single-chain prediction can recover is bounded by the stoichiometry of the native complex, so each positive control was assigned to one of three classes describing how the factor engages DNA at its reference site: monomer, dimer, or larger complex. Classes were assigned from the known DNA-binding stoichiometry of each DBD family, with individual reassignments where the literature on the human ortholog indicates a different assembly. Assignments use only family membership and published stoichiometry, and no prediction-derived quantity. Obligate homo- and heterodimers, including the bZIP, bHLH, and STAT families, were assigned to dimer. Assemblies requiring three or more chains at a single site, such as NF-Y, were assigned to larger complex. Factors with no evidence of obligate oligomerization at the reference site were assigned to monomer. This yielded 139 monomers, 61 dimers, and 10 larger complexes.

### Reconstructing PPMs for multimeric protein complexes

DNA for all multi-chain complexes was drawn from the existing monomeric predictions for each TF. For each TF, folding runs were deduplicated to unique DNA sequences (keeping the highest-scoring run per sequence) and ranked by decreasing *S*_conf_. The DNA from the top 50 scoring structures was carried forward. For complexes of two distinct TFs, the DNA set was the union of both subunits’ top-50 lists. For the Myod1–Tcf3a heterodimer (Fig. 3D; 88 unique sites after removing sequences shared between the two lists), the monomer DNA was used as-is because it already spans the full-length motif.

For the Tp63 and Tp73 homodimers (Supp. Fig. 10A,B; 50 sites each), each monomer site was trimmed to its informative core, the shortest window capturing at least 90% of the IC of its NA-MPNN PPM (relative to a uniform background; minimum 3 bp), and joined to its reverse complement, revcomp(core) + core, reproducing the inverted-repeat geometry of a p53-family half-site from the predicted profile rather than a known consensus, with sequences ranging from 6–20 bp. For the Tp63–Tp73 tetramer, every distinct homodimer sequence *H* was duplicated in tandem without a spacer, *H* + *H*, as in canonical p53-family motifs (12–40 bp).

Each protein chain was taken from that TF’s monomer job, reusing its sequence, precomputed MSAs, and structural templates, so monomer and multimer predictions differ only in stoichiometry and DNA input. NA-MPNN PPMs were predicted as for monomers. Because multimer pools are much smaller than monomer pools, consensus modes for each complex and its matched monomers were built from the top 24 PPMs at a similarity threshold of 0.60, in place of the database setting (47; 0.73), and compared against their JASPAR orthologs.

### Validating TF and motif clusters

To test whether our clustering generalizes across TFs and their motifs, we selected a few representative clusters from the largest DBD families, zf-C2H2 and homeobox. Every member of each selected cluster was co-folded against DNA sequences from HOMER-derived motifs whose BestGuess match belongs to that cluster’s family. PPMs were predicted as in Predicting motifs for all zebrafish TFs and compared to the family motifs both visually and by motif similarity (Supp. Fig. 11A-B). To test whether HOMER clustered motifs correctly, we ran Usf1 (bHLH) against every member of one HOMER motif cluster, 172 motifs that HOMER grouped by similarity (Extracting candidate DNA sequences from Zebrahub-Multiome snATAC-seq), and compared PPM similarity across all reconstructed consensus PPMs, one per Usf1–motif pair (Supp. Fig. 11C).

### Motif-accessibility and TF expression coupling

We used the Zebrahub-Multiome dataset, with paired snATAC + snRNA across 10–24 hpf, integrated into a single Signac object (640,834 peaks; log-normalized RNA). To reduce dropout and stabilize the RNA side of each correlation, cells were aggregated onto the dataset’s precomputed SEACells partition (*~*75 cells per metacell), collating perdataset assignments onto integrated barcodes by (dataset, 10x barcode). Within each metacell ATAC counts were summed and RNA averaged; metacells of fewer than 20 cells were dropped, leaving 1,232 (Supp. Fig. 12A).

ZebraMotif PPMs (the primary binding mode of each TF) and CisBP zebrafish PPMs (one per TF, the motif inferred from the source TF with the highest similarity-regression score) were scanned against the peak set with motifmatchr matchMotifs (dan-Rer11/BSgenome.Drerio.UCSC.danRer11), calling a hit at each motif’s analytic per-motif p-value threshold (*p* < 5 × 10^−5^) (Supp. Fig. 12A). Peaks were harmonized to UCSC seqlevels and clipped to chromosome bounds, and motifs matching ≥1 peak were retained; the resulting differences in match count across motifs are absorbed downstream by chrom-VAR’s GC- and accessibility-matched background correction. Per-motif, per-metacell accessibility deviations were computed on the metacell profiles with chromVAR (getBackgroundPeaks, 50 iterations; computeDeviations), using genome GC bias and non-overlapping peak filtering (Supp. Fig. 12A).

Each PPM was mapped to its TF(s) by stable Ensembl gene ID. Testing was restricted to sequence-specific TFs on a curated AnimalTFDB4 whitelist that were expressed in ≥5% of metacells and whose PPM carried ≥1 bit of total IC, since near-zero-IC motifs match tens of thousands of peaks and self-rank spuriously. For each motif we computed the Spearman correlation between its deviation z-score and its own TF’s expression across metacells, then ranked this own-TF correlation against the correlations of the same deviation track with all other pooled TFs. Same-family paralogs were excluded because they share the motif. This yielded an own-TF percentile (null median 0.5) (Supp. Fig. 12B). Because CisBP is small and curated and ZebraMotif covers 2,543 TFs and is dominated by zf-C2H2, we compared the two databases within DBD classes so that coupling is not confounded by family composition. Classes reconcile the family vocabularies of AnimalTFDB4, CisBP and JAS-PAR (TFClass) through assignment based on Inter-Pro DBD accessions. For example, the zf-C2H2 class merges the AnimalTFDB4 zf-C2H2 and ZBTB families (Supp. Fig. 12B; Fig. 4D).

### Gene regulatory network inference w ith CellOracle

To test whether ZebraMotif PPMs recover regulatory biology, we ran CellOracle^36^ on Zebrahub-Multiome with two motif sets for the base gene regulatory network (GRN): CisBP alone, and a hybrid in which every TF retains both its CisBP and ZebraMotif motifs. All other parameters were identical, so GRN differences reflect the motif database alone.

For each of six timepoints (0, 5, 10, 15, 20, and 30 somites), peaks linked to promoters by Cicero co-accessibility were scanned with each motif set on danRer11 (FPR 0.02, score threshold 10), admitting a TF→target edge wherever a TF motif hit a peak linked to the target gene (Supp. Fig. 13A). Base GRNs were fit to the matched snRNA-seq following the standard CellOracle workflow. Per timepoint, the top 3,000 highly variable genes were KNN-imputed after PCA, and within each of 32 cell types each target gene was regressed on its candidate regulators by bagging ridge regression (α = 10). Coefficients give edge sign and strength; edges with *p* < 0.001 were retained, up to 2,000 per cell type, giving 192 cell-type × timepoint GRNs per motif set.

For ego-regulons (Supp. Fig. 13B), we selected four (*tbx6, her7, tox3, prdm1a*) of the 21 TFs whose expression–accessibility coupling was detected with ZebraMotif but not CisBP (own-TF percentile >95th with strong cell-type correlation; Motif-accessibility and TF expression coupling) and plotted their top 12 significant targets by absolute coefficient in the hybrid GRN. The PSM network (Fig. 4E) was built identically from the CisBP and hybrid 15-somite PSM GRNs: the top eight significant edges per seed (*tbx6, tbx16, her1, her7*) by absolute coefficient, plus all significant edges among the selected nodes, drawn on a shared layout.

### Compute

The screen consumed approximately 107,000 GPU-hours at a mean of roughly three minutes per job (25 predictions per job). On a sample of 100,000 jobs, 55% ran on H200 or H100 GPUs at 2–2.25 minutes per job, 30% on A40 or A6000 GPUs at 4.5–4.75 minutes, and the remaining 15% on A100 or L40S GPUs at 3–3.5 minutes.

---

**Supplementary Fig. 1.**
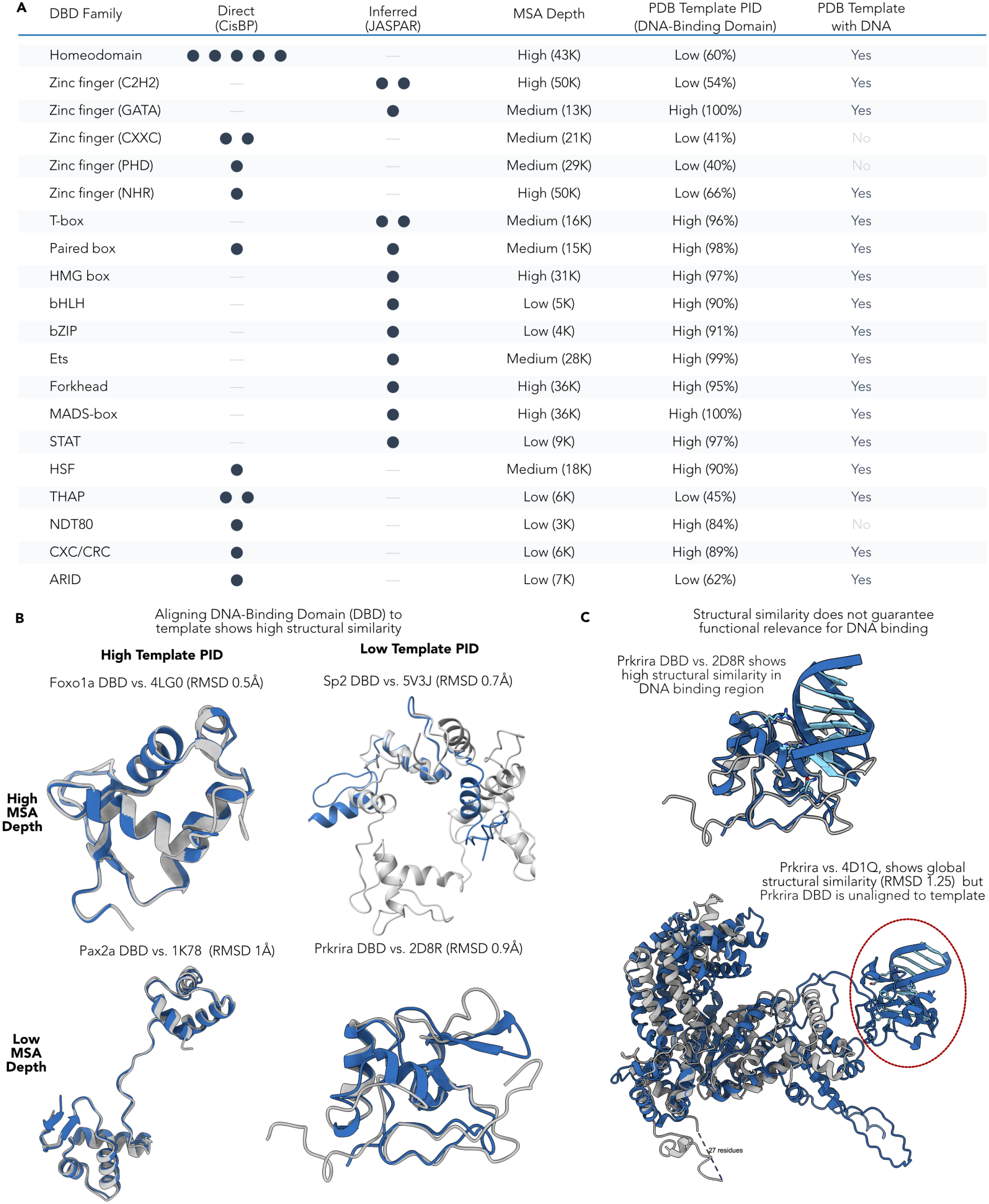
Selecting TFs for motif–non-motif experiments. **(A)** Benchmark TFs span DBD families, MSA depths, and template availability. Across 20 DNA-binding domain (DBD) families, 16 TFs were taken from CisBP and 13 from JASPAR. MSA depth is binned as Low (<10K sequences), Medium (10K–30K), or High (≥30K) to observe if failures track sequence depth. DBD sequence identity to the templates returned by the PDB search is binned as Low (<70%) or High (≥70%), and TFs with no returned template are marked, to test whether accurate structural templates exist for each DBD. (**B)** Predicted DBDs superpose closely on their PDB templates regardless of MSA depth or template identity. Example DBDs aligned to their closest templates, with backbone RMSD, at high and low MSA depth and template percent identity (PID). **(C)** Global fold accuracy does not guarantee correct DBD placement. For Prkrira, aligning only the DNA-bound DBD to the template gives high structural similarity. Aligning the entire predicted structure also gives high global similarity, but leaves the DBD not aligned to any part of the template. The template search can therefore return templates that match a protein at one level of organization but not another.

**Supplementary Fig. 2.**
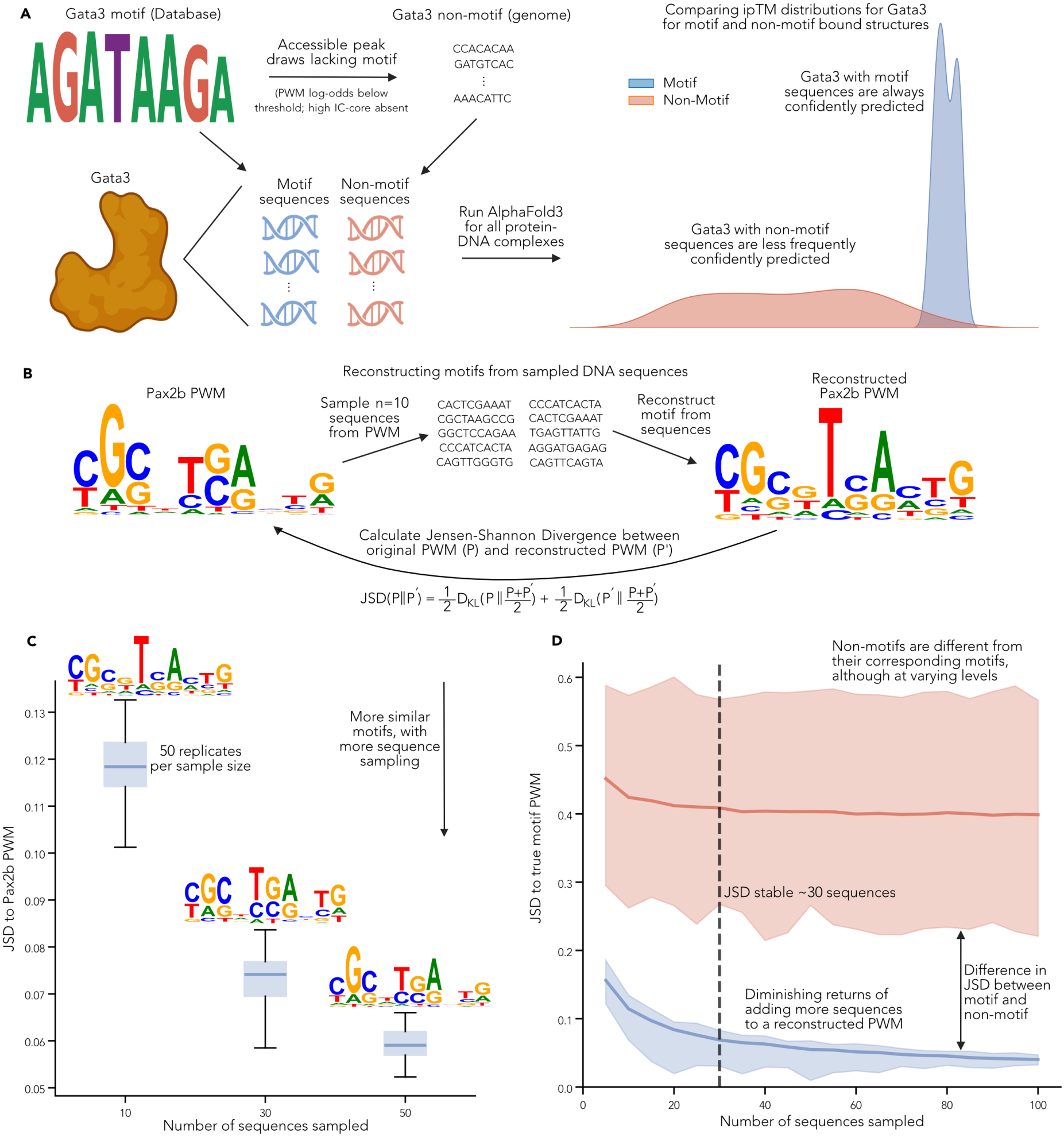
Designing motif–non-motif experiments. **(A**) Motif–non-motif design, illustrated for Gata3. Motif sequences are sampled from the orthologous JASPAR motif, and GC- and length-matched non-motif sequences are drawn from the genome. Gata3 is co-folded with each DNA sequence individually, and confidence distributions are compared between motif- and non-motif-bound structures. **(B)** PWM reconstruction is measured by Jensen–Shannon divergence (JSD). Sequences are sampled from a PWM (here *n* = 10 from Pax2b), a PWM is reconstructed from per-position frequencies, and JSD to the source PWM is computed. **(C)** JSD falls with sample size. Repeating (B) with increasing numbers of sampled sequences, JSD to the Pax2b source PWM decreases up to 50 sequences. **(D)** Thirty sequences per PWM is enough to reconstruct PWMs for most TFs. Across all 29 benchmark TFs, JSD levels off near 30 sampled sequences for most reconstructed PWMs. PWMs reconstructed from non-motif genomic controls remain roughly stable across sample sizes, decreasing only slightly as more sequences are sampled.

**Supplementary Fig. 3.**
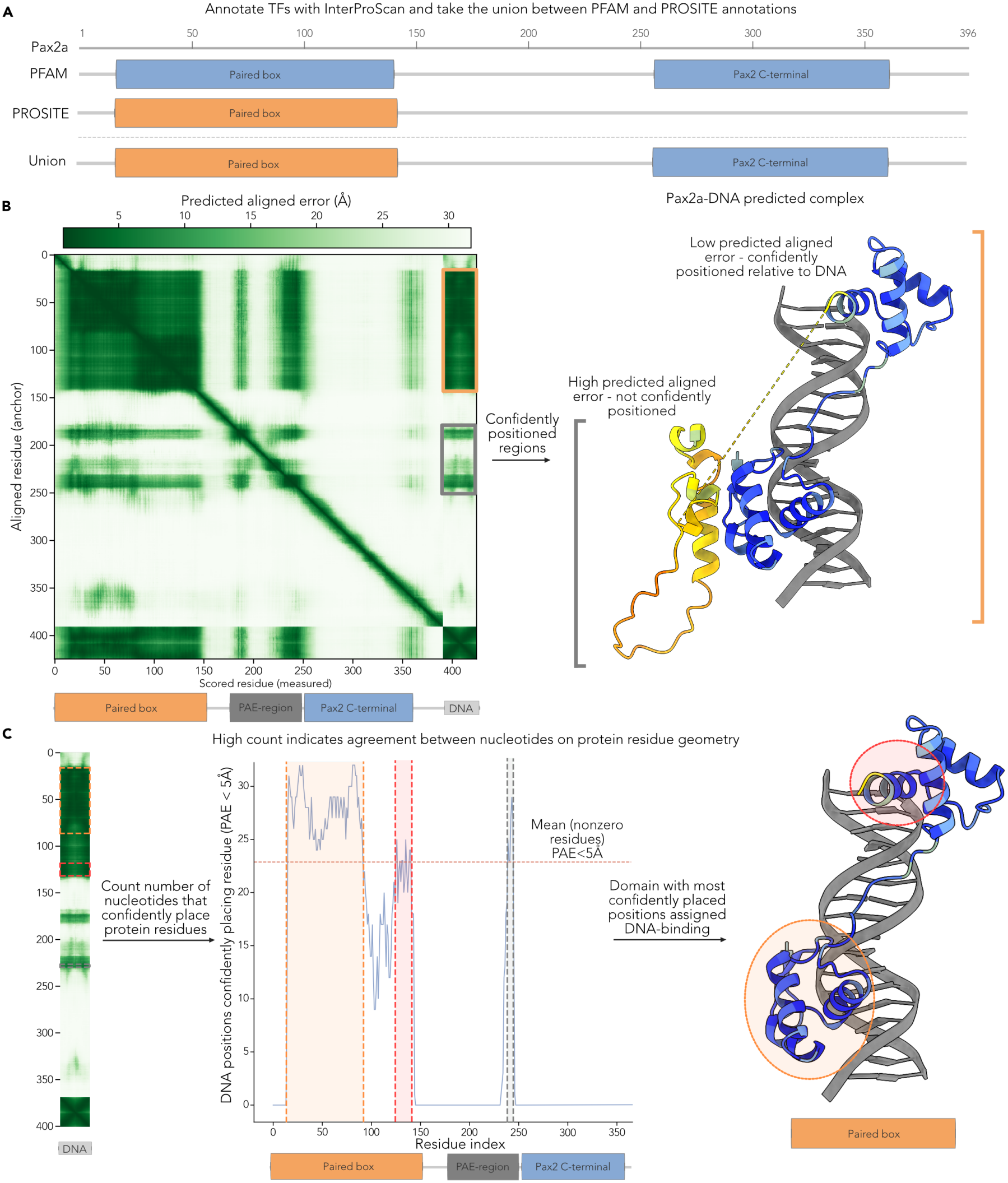
The DNA-binding domain (DBD) is called as the annotated domain that AF3 most confidently positions against the DNA. **(A)** Domains are annotated with InterProScan as the union of Pfam and PROSITE calls, with Pax2a shown as an example. **(B)** AF3 confidently positions the paired box, but not the C-terminal region, relative to the DNA. Low PAE between paired-box residues and the DNA (orange box) indicates that AF3 is confident in their relative position; the Pax2 C-terminal region has high PAE to the DNA. Low PAE reflects confident relative positioning, not necessarily physical contact. **(C)** PAE localizes the DBD to the paired box. For each residue, we count the DNA nucleotides placed with PAE < 5 Å and retain residues at or above the mean nonzero count (dashed line). The annotated domain with the highest mean frequency of retained residues across predictions, here the paired box, is assigned as the DBD.

**Supplementary Fig. 4.**
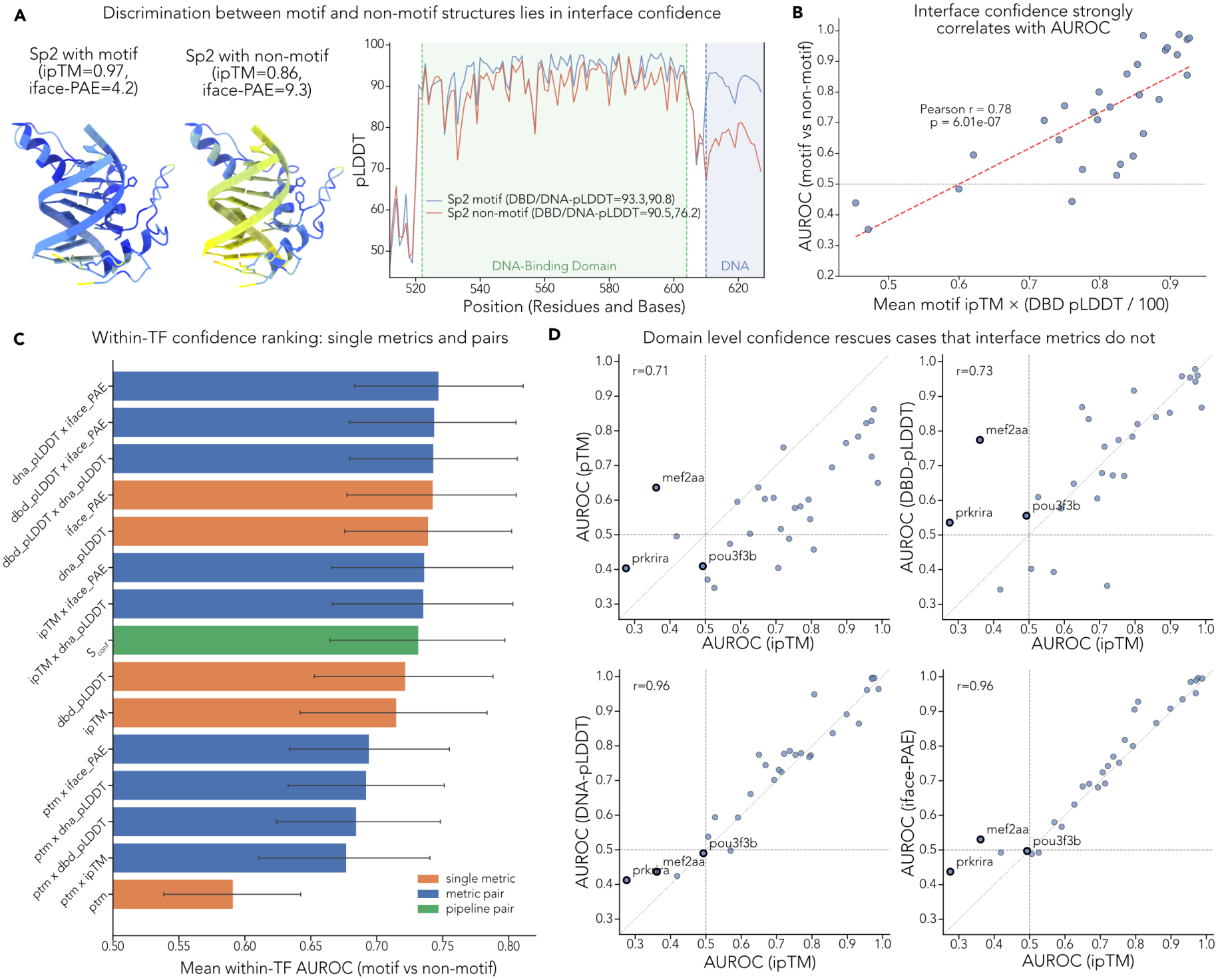
Selecting the appropriate confidence metrics to determine interface confidence. **(A)** Interface confidence metrics, such as ipTM and iface-PAE which are computed by AF3, and pLDDT averaged across the DNA-binding domain (DBD-pLDDT), show higher confidence in the Sp2 motif-bound structure than the Sp2 non-motif bound structure. **(B)** Interface confidence, measured by the mean ipTM x (DBD pLDDT 100) of all motif-bound structures per-TF, correlates strongly with AUROC across all 29 TFs in the motif–non-motif panel. **(C)** No interface metric or pairwise combination discriminates substantially better than the others. The mean within-TF AUROC for each interface confidence metric, and combination of metrics, all fall within overlapping confidence intervals. pTM, a global confidence metric, is shown for comparison and discriminates less well than interface-specific metrics. **(D)** DBD-pLDDT recovers TFs that the other metrics miss. Per-TF AUROC of pTM, DBD-pLDDT, DNA-pLDDT, and interface PAE, each plotted against ipTM. DBD-pLDDT is the only metric that discriminates both Mef2aa and Pou3f3b above chance, so ipTM and DBD-pLDDT are combined into *S*_conf_.

**Supplementary Fig. 5.**
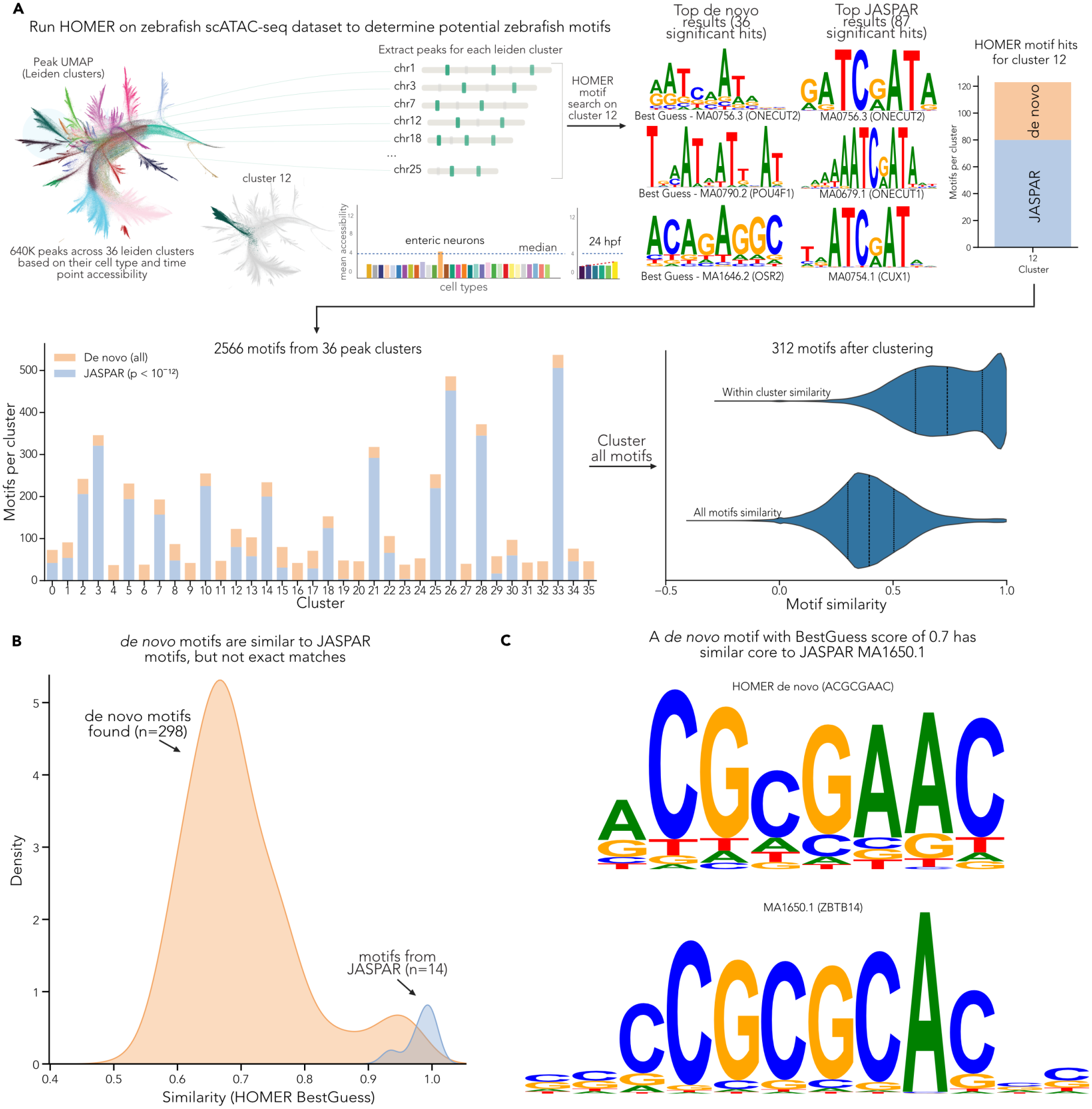
Discovering motifs from accessible chromatin in Zebrahub-Multiome data. **(A)** Motifs are discovered separately within each regulatory program. HOMER is run on all peaks within each of the 36 Leiden clusters that group over 640,000 peaks based on their cell type and time point accessibility in 10-24 hours-post-fertilization (hpf) developing zebrafish. Each cluster is enriched for particular cell types and time points that give rise to motifs corresponding to the developmental program in each cluster. For example, cluster 12 is enriched for enteric neurons with accessibility rising from 10 to 24 hpf. HOMER returns *de novo* motifs, all of which are retained, and JASPAR matches, retained at *p* < 10^−12^ (cluster 12: 43 and 80, respectively). Across all clusters this gives 2,566 motifs, which reduce to 312 non-redundant motifs after clustering. **(B)** *De novo* motifs resemble JASPAR motifs without duplicating them. BestGuess score (HOMER’s similarity to the closest JASPAR match) of each of the 298 *de novo* motifs (median 0.68). The remaining 14 library motifs are JASPAR matrices from HOMER’s known-motif enrichment. **(C)** A *de novo* motif at the median BestGuess score shares its core with its closest JASPAR match. The closest JASPAR match (BestGuess) of this *de novo* motif is MA1650.1, assigned to human ZBTB14. Both cores read similar consensus sequences (CGCGAAC and CGCGCA for the *de novo* and JASPAR motifs, respectively), but information content differs within the core.

**Supplementary Fig. 6.**
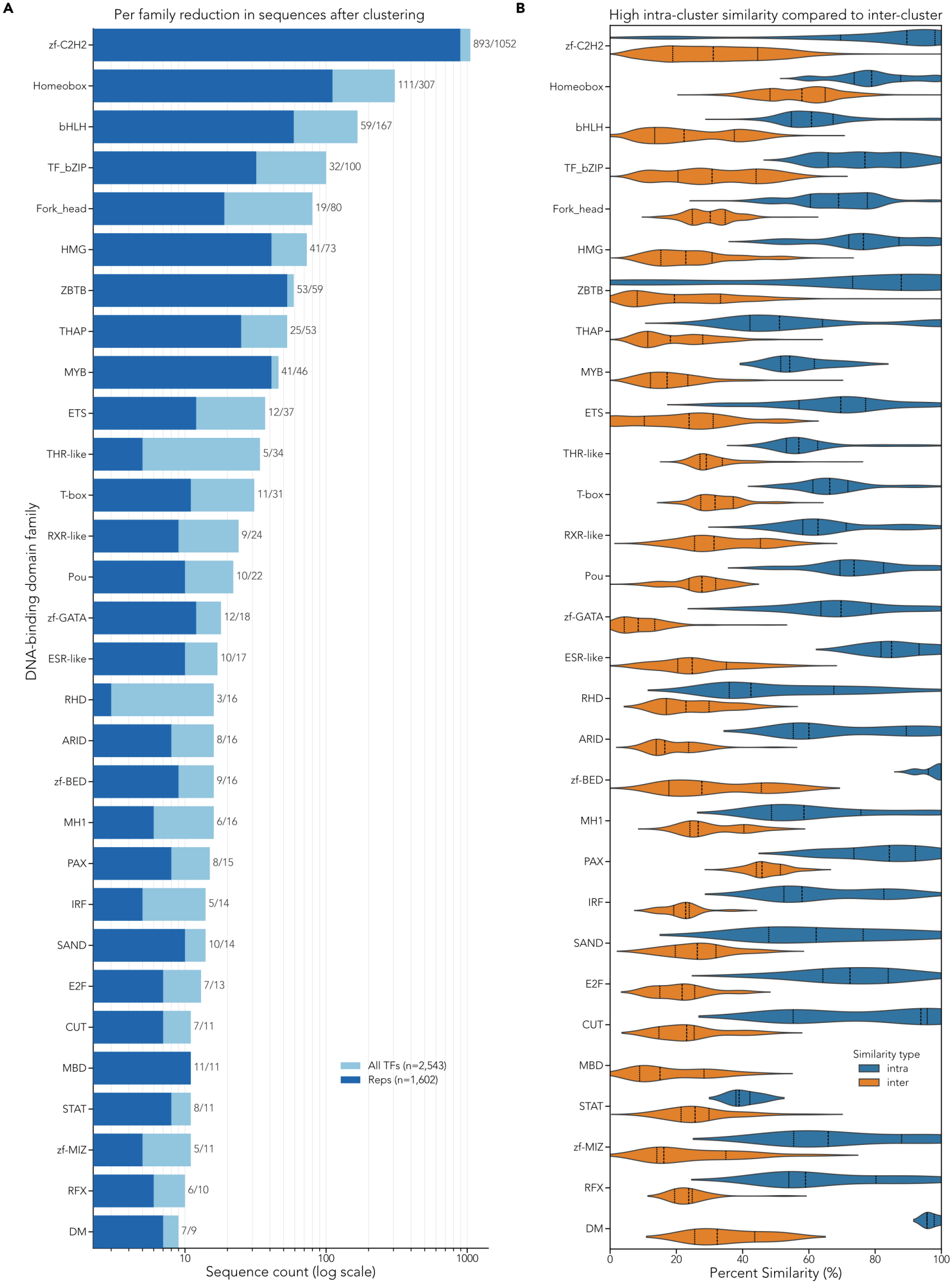
Clustering TFs reduces counts in nearly all families and maintains similarity w ithin families. **(A)** Nearly all TF families (showing 30 of 72, ranked by number of members) are reduced by clustering, with zf-C2H2 and ZBTB being among the least reduced due to the strict recognition code procedure we apply to ensure similarity is considered across large, multi-domain finger arrays. **(B)** Members of a cluster are more similar to one another than to members of other clusters. Sequences are aligned within each cluster and across each pair of clusters to compute intra- and inter-cluster percent similarity.Across families, intra-cluster similarity exceeds inter-cluster similarity.

**Supplementary Fig. 7.**
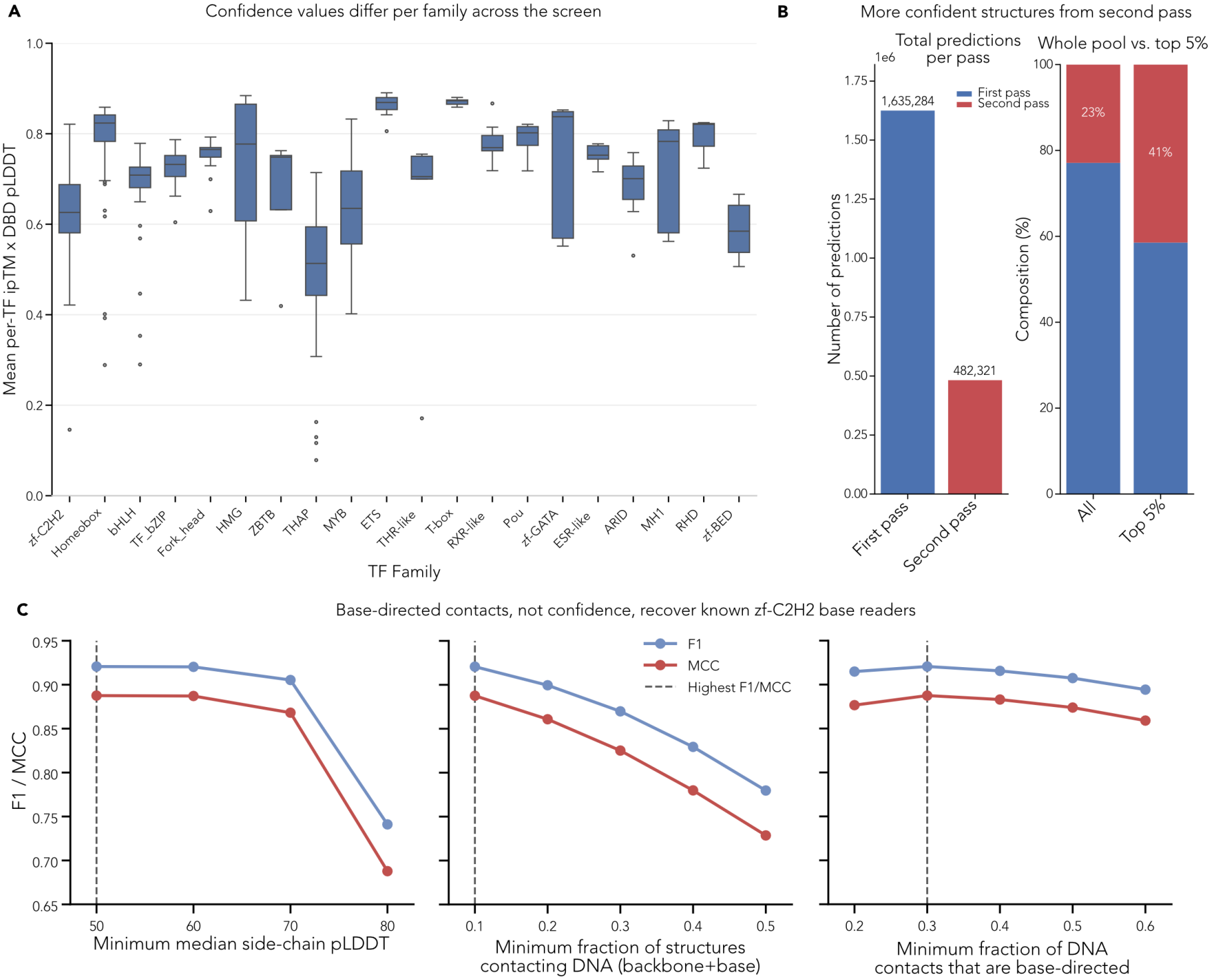
Screen confidence across DNA-binding domain (DBD) families, second-pass enrichment, and calibration of base-reader calls. **(A)** Confidence is family-dependent but high for most TFs. Each box summarizes per-TF mean ipTM × DBD-pLDDT across all of that TF’s predicted structures, for the 20 largest DBD families ordered by size. Most families center between 0.7 and 0.9 with tight spread; THAP, zf-BED, zf-C2H2, and MYB sit lower, and HMG and zf-GATA are broadly spread. **(B)**The second pass concentrates confident structures. Left, structures predicted in the first (1,635,284) and second (482,321) passes. Right, pass composition of the whole pool versus the top 5% of structures by *S*_conf_: the second pass contributes 23% of all structures but 41% of the top 5%. **(C)** The base-directed fraction of contacts identifies known base-reading residues in zf-C2H2 TFs. A residue is called a base reader when it passes three gates: median side-chain pLDDT exceeds a minimum (left); its side chain lies within 5 Å of DNA in at least a minimum fraction of the TF’s structures (middle); and at least a minimum fraction of those contacts are to bases rather than the backbone (right). Calls were scored against recognition-code positions p−1, p2, p3, and p6 within the DNA-binding fingers of 756 zf-C2H2 TFs (8,072 positives); negatives are the remaining finger residues. Each subpanel varies one gate with the other two held at the selected operating point (dashed line; pLDDT 50, contact fraction 0.1, base fraction 0.3; F1 0.921, MCC 0.88).Raising the pLDDT gate above 70 or the contact-frequency gate above 0.1 lowers F1 and MCC by discarding true readers while precision stays near 0.97, so neither gate adds specificity. The base-fraction gate is the only one that shifts the precision–recall balance, and it is nearly flat around its optimum. Residues that AF3 places against bases in as few as 10% of structures are therefore overwhelmingly those the recognition code predicts.

**Supplementary Fig. 8.**
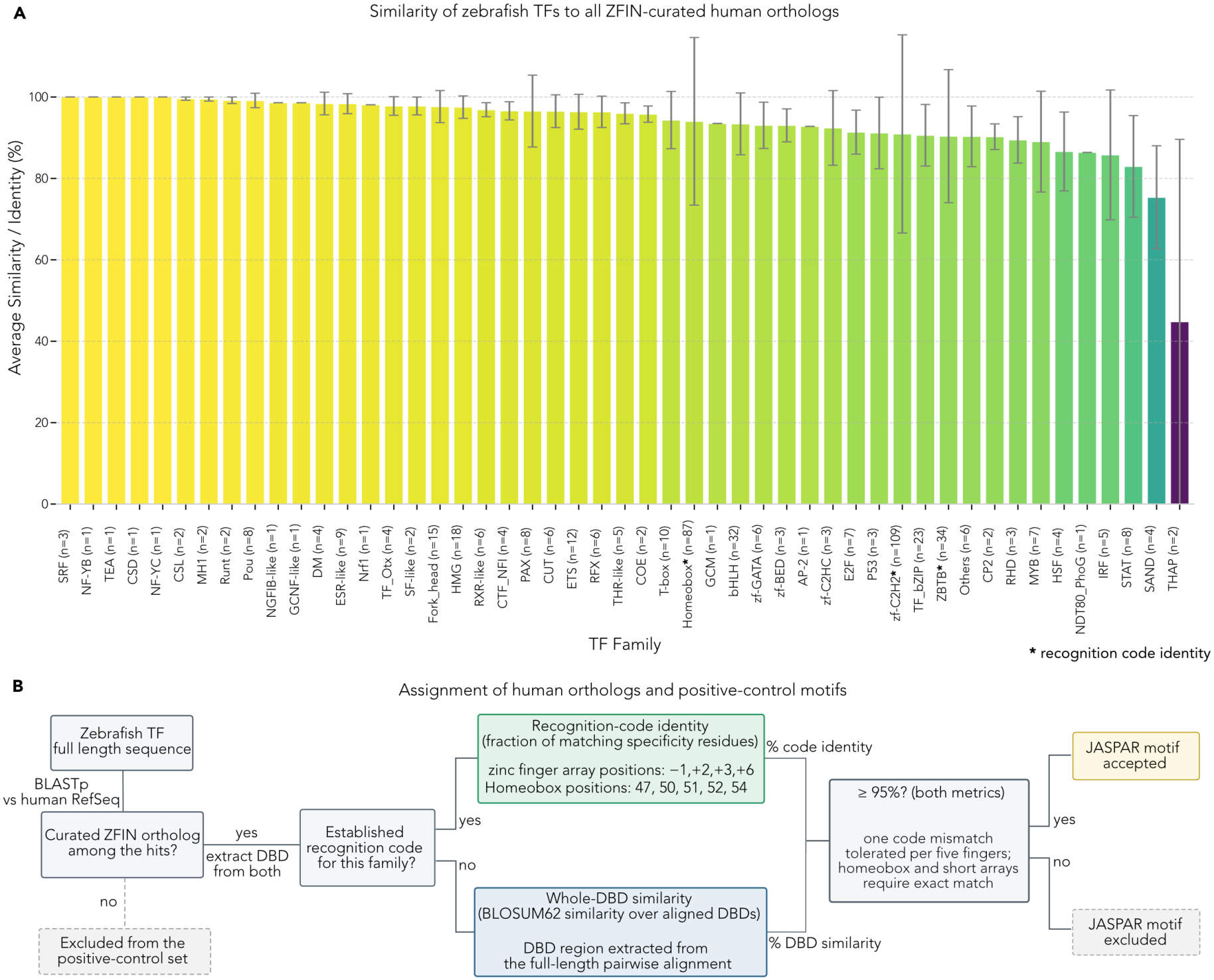
Determining similarity between representative zebrafish TFs and their human orthologs w ith JASPAR motifs. **(A)** Most DNA-binding domain (DBD) families retain >85% similarity to their human orthologs. Bars show the per-family mean of DBD similarity (BLOSUM62; unmarked) or recognition-code identity (asterisk; zf-C2H2, ZBTB, Homeobox) between each zebrafish TF and its ZFIN-curated human ortholog, ordered from highest to lowest; error bars, s.d.; 486 TFs. Recognition-code families show the widest spread because code scoring is coarse: a substitution at p−1 or p3 zeroes the whole finger, and a single homeodomain substitution costs 20%. **(B)** Workflow for assigning positive-control motifs. Each zebrafish TF’s full-length sequence is searched against human RefSeq with BLASTP, and TFs whose ZFIN-curated ortholog is absent from the hits are excluded. DBDs are extracted from both proteins and compared by recognition-code identity for families with an established code, or by BLOSUM62 similarity over the aligned DBDs. The ortholog’s JASPAR motif is accepted as ground truth when the score reaches ≥95%. This yields 210 positive controls (Positive-control benchmark and validation of predicted PPMs).

**Supplementary Fig. 9.**
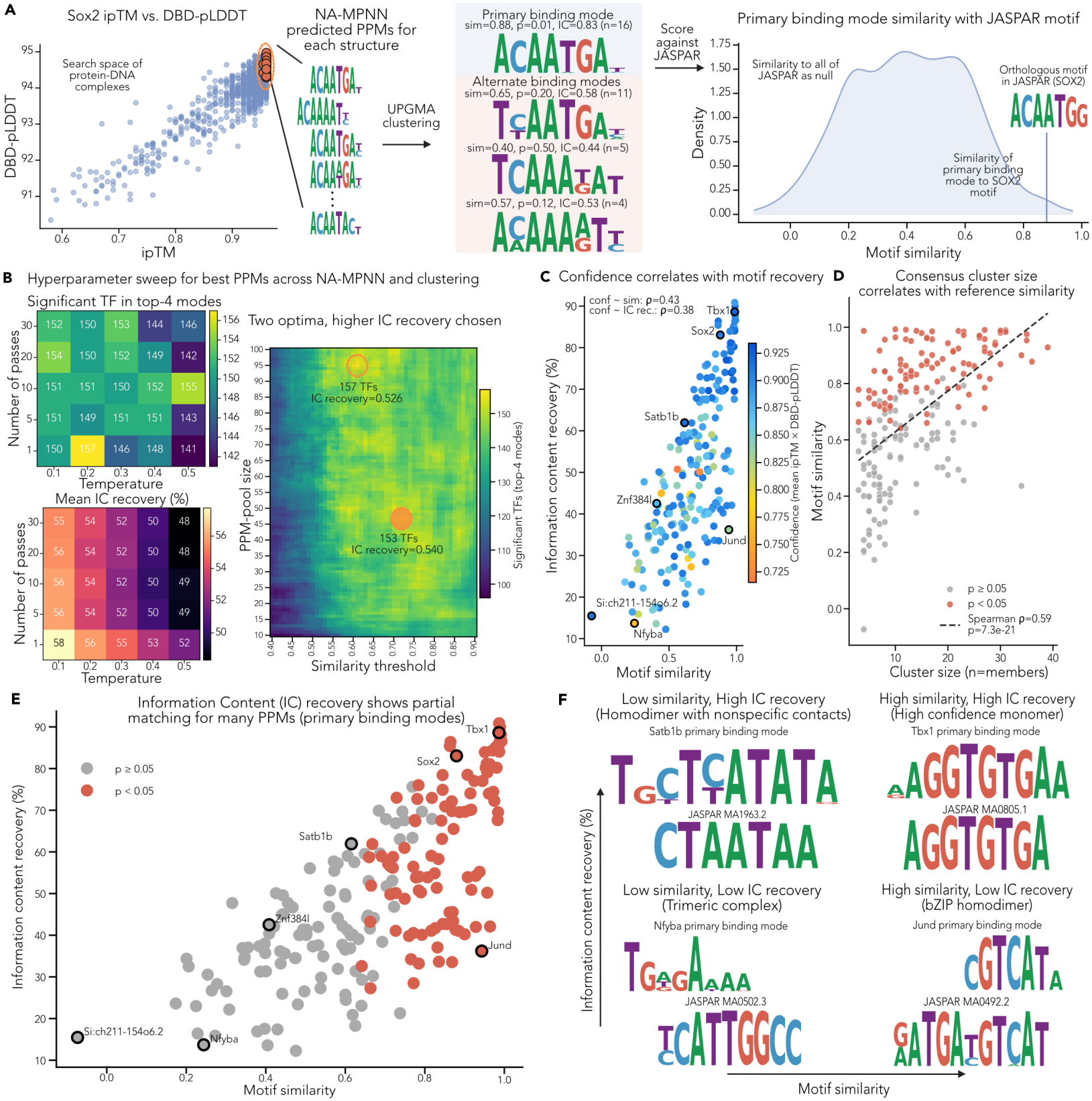
Selecting PPM-prediction hy perparameters on the positive-control set and characterizing recovery. **(A)** Worked example for Sox2. Predicted complexes span a range of ipTM and DBD-pLDDT values. The most confident are passed to NA-MPNN, which predicts one PPM per structure, and UPGMA clustering condenses these into a primary binding mode (16 PPMs; motif similarity 0.88, *p* = 0.01, information content (IC) recovery 0.83) and alternate modes. The primary mode is scored against the orthologous SOX2 motif with all of JASPAR as the null distribution. **(B)** Hyperparameter sweep. Left: positive controls with a significant JASPAR match within the top four modes (top) and mean information content (IC) recovery (bottom) for each NA-MPNN temperature and number of passes per structure. Each cell is the best value over a coarse consensus grid (PPM-pool size 10–30), with the two metrics maximized independently. IC recovery decreases with temperature at every pass count, while significant-TF counts vary little. Right: significant TFs across PPM-pool size (10–100) and similarity threshold for the selected NA-MPNN setting (temperature 0.1, 30 passes). Of two similar optima (157 TFs at IC recovery 0.526; 153 TFs at 0.540), the one with higher IC recovery was deployed (47 PPMs, threshold 0.73). **(C)** Confidence stratifies quality within the high-confidence structures. IC recovery versus motif similarity to the orthologous motif, colored by mean ipTM × DBD-pLDDT (Spearman *ρ* = 0.43 with similarity and 0.38 with IC recovery; *n* = 210). **(D)** Agreement among predicted PPMs is a stronger indicator of a matched motif than structure confidence. Motif similarity versus the number of PPMs in the primary mode (Spearman *ρ* = 0.59, *p* = 7.3 × 10^−21^). **(E)** IC recovery reveals partial matches among non-significant modes. IC recovery versus motif similarity for primary modes, colored by significance. **(F)** Examples across the similarity–IC space. Tbx1, a high-confidence monomer, matches on both. Satb1b, a homodimer with nonspecific contacts, reaches high IC recovery at low similarity through AT-rich positions. Jund, a bZIP homodimer, matches one half-site of its full reference. Nfyba, a non-specific subunit of the trimeric NF-Y complex, matches no part of its reference.

**Supplementary Fig. 10.**
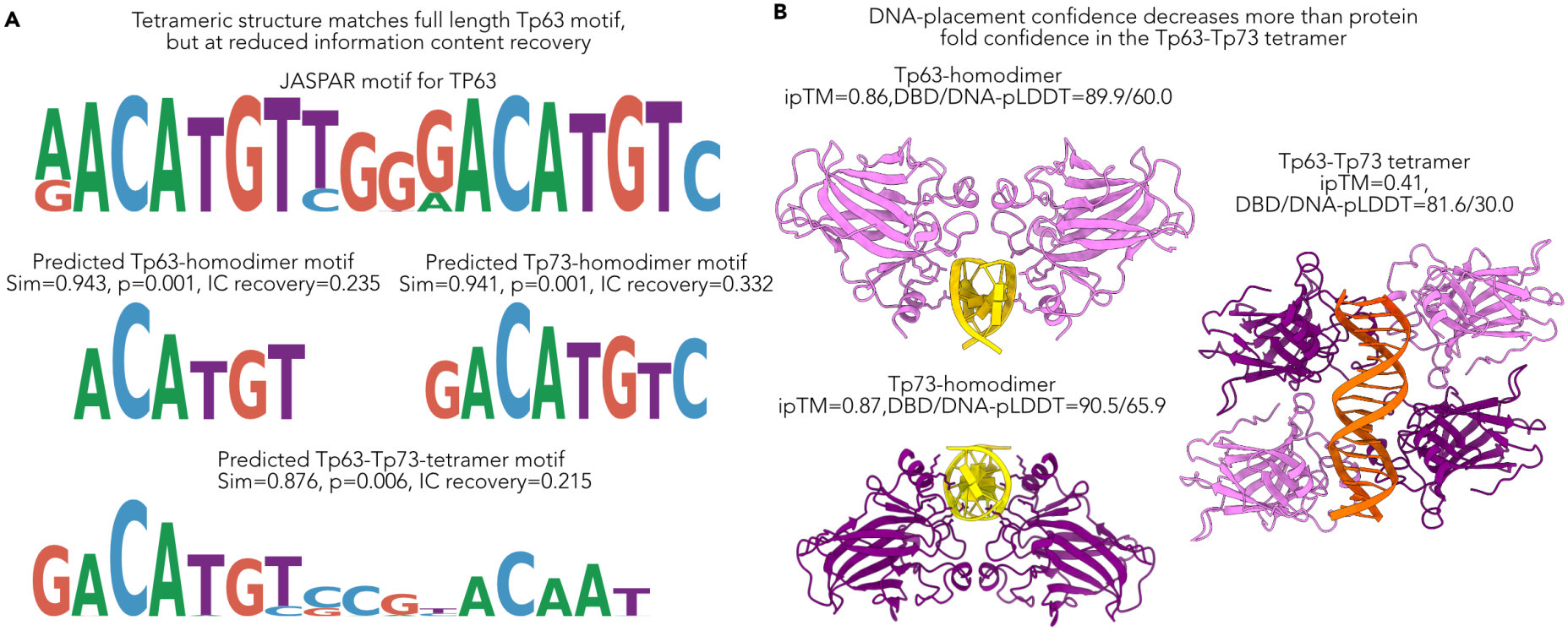
Modeling a Tp63–Tp73 tetramer spans the full p53-family motif but does not improve motif recovery. **(A)** The tetramer recovers the length of the TP63 motif, but with lower motif similarity and information content (IC) recovery than either homodimer achieves from a single half-site. Top, the full-length JASPAR TP63 motif. Below, primary binding modes predicted for the Tp63 homodimer (motif similarity 0.943, *p* = 0.001, IC recovery 23.5%) and the Tp73 homodimer (0.941, *p* = 0.001, 33.2%), each covering one half-site centered on the CATG core, and for the Tp63–Tp73 tetramer (0.876, *p* = 0.006, 21.5%), which pairs a well-predicted first half-site with a weakly predicted second. **(B)** The tetramer loses confidence mainly in DNA placement. Representative structures of the Tp63 homodimer (ipTM 0.86; DBD/DNA-pLDDT 89.9/60.0), the Tp73 homodimer (0.87; 90.5/65.9), and the 2:2 Tp63–Tp73 tetramer (0.41; 81.6/30.0). Tp63, pink; Tp73, purple; DNA, medium confidence (homodimers) and low confidence (tetramer).

**Supplementary Fig. 11.**
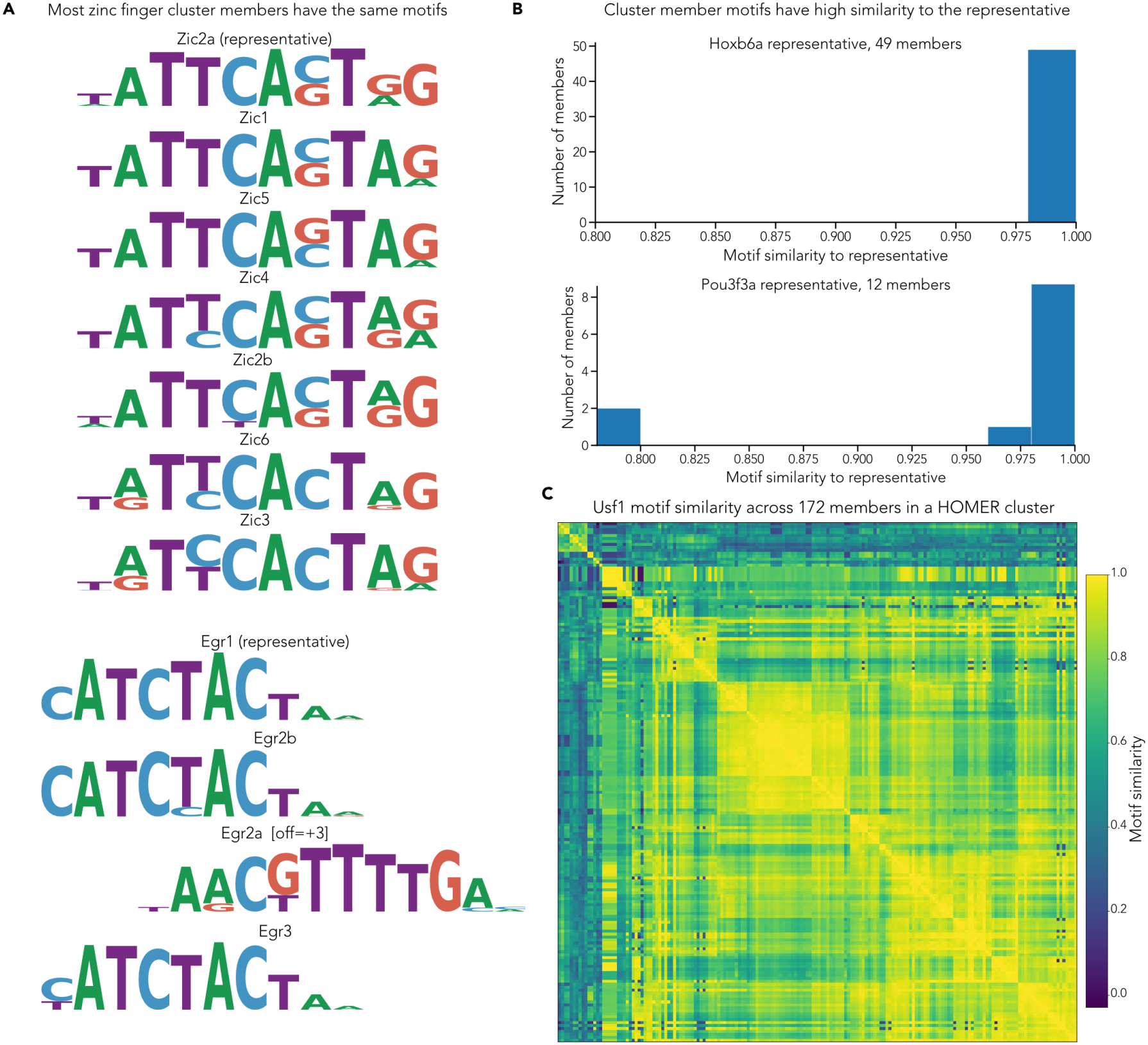
Most cluster members predict PPMs similar to their representative’s. **(A)** Members of zf-C2H2 recognition-code clusters reproduce the representative’s primary binding mode. Every member of the Zic and Egr clusters was co-folded with the same three HOMER motif clusters, so members differ only in protein sequence. All six Zic members reproduce the Zic2a primary mode; Egr2b and Egr3 reproduce the Egr1 prediction, and Egr2a predicts a distinct motif (shown aligned to the representative at a +3 bp offset). (B)Homeodomain cluster members predict PPMs nearly identical to the representative’s. Histograms show the motif similarity of each member’s predicted PPM to the representative’s. All 49 members of the Hoxb6a cluster score above 0.975, and 10 of 12 members of the Pou3f3a cluster score above 0.96; the remaining two fall near 0.80. **(C)**Motifs grouped by HOMER drive a common predicted specificity. Pairwise motif similarity between the consensus PPMs that Usf1 (bHLH) predicts from each of 172 motifs in a single HOMER motif cluster, with rows and columns ordered by hierarchical clustering. Similarity is high across most of the matrix, supporting the use of one representative motif per HOMER cluster; a small block of motifs (top left) is dissimilar to the rest.

**Supplementary Fig. 12.**
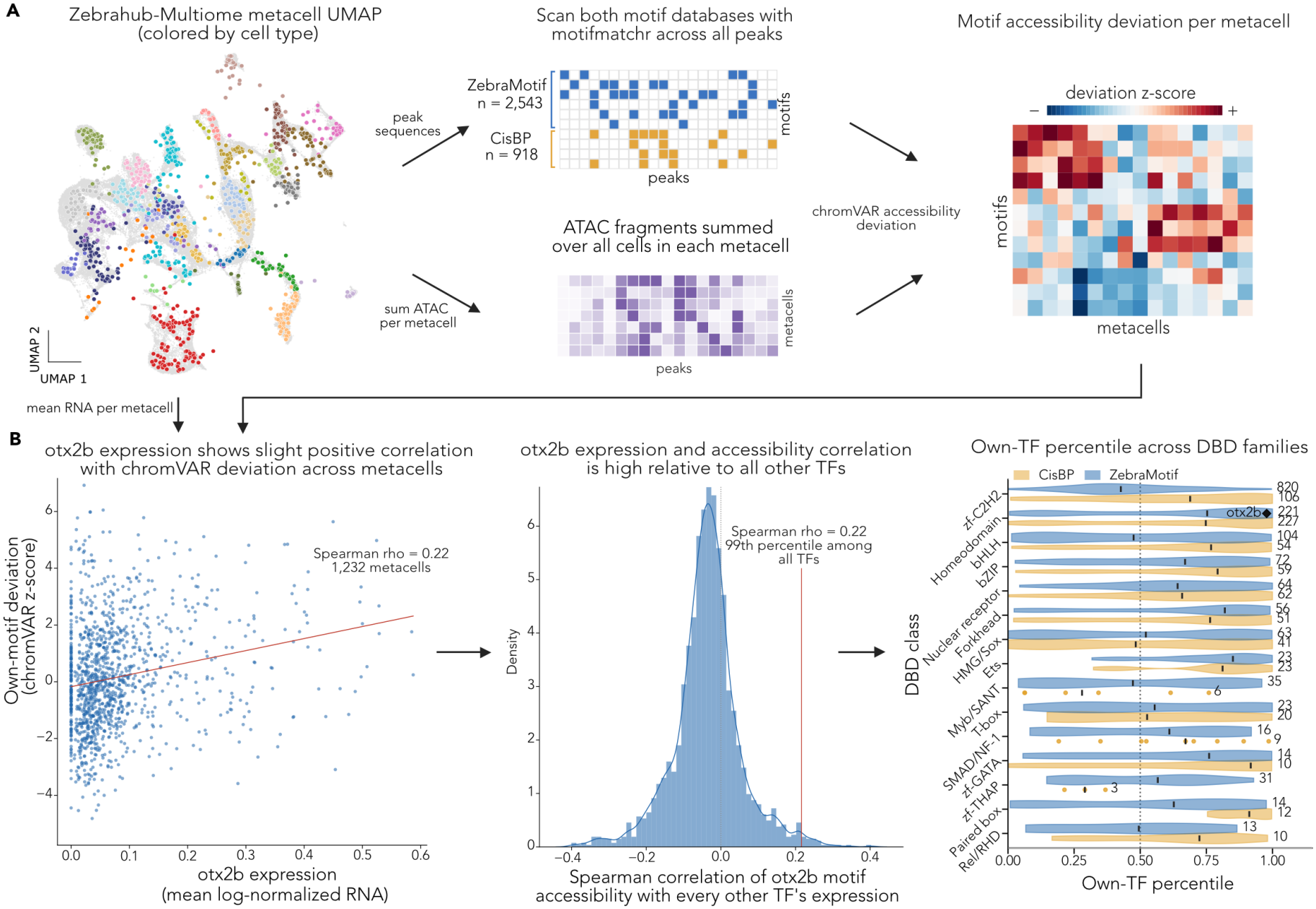
Measuring TF expression–motif accessibility coupling w ith chromVAR. **(A)** chromVAR scores each motif’s accessibility in every metacell from two independent inputs. Zebrahub-Multiome cells are aggregated into metacells (UMAP, colored by cell type). Motif hits are called from peak sequences by scanning ZebraMotif (*n* = 2,543) and CisBP (*n* = 918) PPMs against all peaks with motifmatchr, while ATAC fragments are summed over the cells in each metacell. chromVAR combines the two into a per-motif, per-metacell accessibility deviation z-score, and mean RNA per metacell supplies TF expression for (B). **(B)** Own-TF coupling ranks each motif’s expression–accessibility correlation against every other TF. Left, *otx2b* expression is modestly correlated with the deviation of its own motif across 1,232 metacells (Spearman *ρ* = 0.22). Middle, correlating the same deviation with the expression of every other TF (same-family paralogs excluded) forms a null against which *otx2b*’s own correlation (red line) falls at the 99th percentile. Right, own-TF percentiles by DNA-binding domain (DBD) family for CisBP (gold) and ZebraMotif (blue), the comparison summarized in Fig. 4D; the diamond marks *otx2b*.

**Supplementary Fig. 13.**
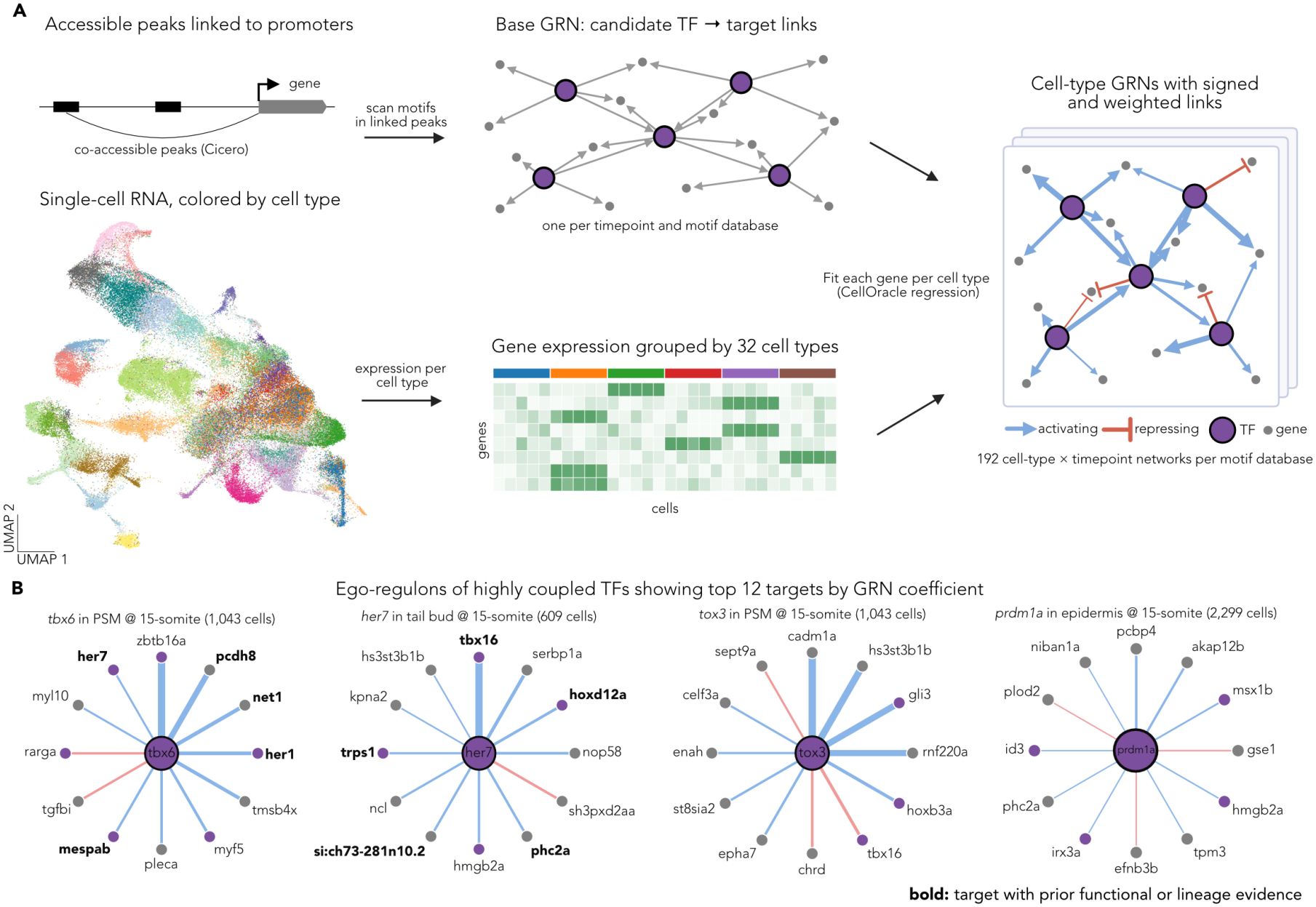
CellOracle gene regulatory network (GRN) inference w ith ZebraMotif motifs. **(A)** Each motif database yields its own set of cell-type-specific, signed GRNs. Accessible peaks linked to gene promoters by Cicero co-accessibility are scanned for TF motifs. This gives a base GRN of candidate TF→target links for each timepoint and motif database. Matched snRNA-seq (UMAP colored by cell type) is grouped into 32 cell types, and CellOracle regression fits each target gene on its candidate regulators within each cell type. The resulting edges are signed (blue, activating; red, repressing) and weighted by regression coefficient, giving 192 cell-type × timepoint networks per motif database. Purple nodes are TFs and grey nodes are target genes. **(B)** The hybrid GRN assigns highly coupled ZebraMotif TFs lineage-consistent candidate targets. Ego-regulons show the top 12 targets by absolute GRN coefficient (*p* < 0.001) for *tbx6* and *tox3* in the presomitic mesoderm (PSM), *her7* in the tail bud, and *prdm1a* in the epidermis, all at the 15-somite stage. Edge width scales with absolute coefficient and edge color gives sign. Bold labels mark targets with prior functional or lineage evidence.

**Supplementary Fig. 14.**
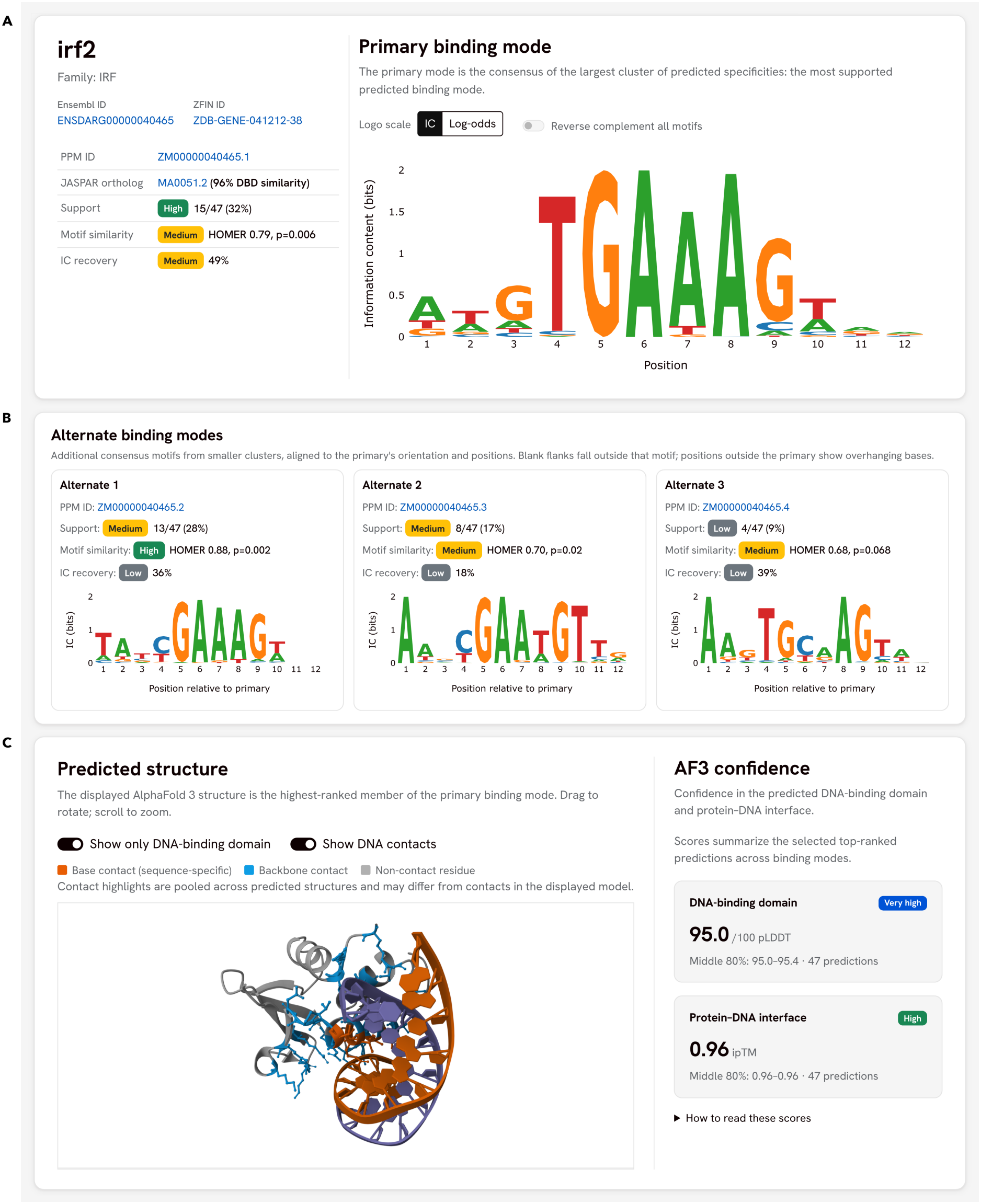
The ZebraMotif web interface. Entry page for Irf2 at zebramotif.apps.czbiohub.org. **(A)** Each entry reports the TF’s primary binding mode, the consensus of the largest cluster of predicted position probability matrices, with its cluster support, similarity to the closest JASPAR ortholog and IC recovery. **(B)** Alternate binding modes from smaller clusters are aligned to the primary mode and have the same annotations. **(C)** The highest-ranked AF3 structure of the primary mode, restricted to the DBD and with residues colored by base or backbone contact pooled across the supporting structures, alongside DBD-pLDDT and ipTM summarized over the 47 top-ranked predictions across all binding modes. A walkthrough of the website is provided as Supplementary Video 1.

**Table 1.** Software, databases, and parameters. Parameters not listed were left at package defaults.

| Resource | Version / build | Key parameters and notes |
|---|---|---|
| *Structure and specificity prediction* |  |  |
| AlphaFold3<sup>19</sup> | v3.0.1-23-g792e61e; Nov. 2024 model parameters | Five model seeds × five samples (25 predictions, all considered one job); top-ranked prediction retained per job. Full-length protein; DBD used only for scoring and downstream analysis. DNA supplied as two entities (forward and reverse complement); PWM-sampled sequences have the width of their source PWM. |
| AF3 sequence databases | Downloaded Nov. 2024 | Small BFD; MGnify 2022_05; UniProt 2021_04 (paired); UniRef90 2022_05; NT-RNA 2023_02_23 (clustered 90/80); Rfam 14.9; RNAcentral (linclust 90/80). Paired and unpaired MSAs from the CPU pipeline passed unmodified to GPU inference. |
| AF3 templates | mmCIF mirror 11 Oct. 2024; PDB seqres 2022_09_28 | Enabled, maximum template release date 30 Sep. 2021 (matching the AF3 publication); no cap on template count. |
| NA-MPNN<sup>24</sup> | commit 835974f3; checkpoint s_70114.pt | Specificity mode on the full AF3 complex (mmCIF → PDB). batch_size = 1, number_of_batches = 30 (30 passes per structure), unfixed sampling seed, temperature 0.1 (Sweeping parameters across PPM prediction space). |
| *Sequence annotation, alignment, and clustering* |  |  |
| InterProScan<sup>43</sup> | 5.73-104.0 | Union of Pfam 37.2 and PROSITE 2023_05 matches (member-database releases as bundled with this InterProScan build); overlapping calls merged into one contiguous span. |
| pyhmmer<sup>45</sup> | 0.12.0 | hmmalign of homeodomains to the Pfam PF00046 HMM.<sup>44</sup> |
| MMseqs2<sup>32</sup> | 18.8cc5c | easy-cluster, defaults, per family on domain-trimmed sequences; zf-C2H2, ZBTB, and Homeobox clustered by recognition code instead (Recognition codes). |
| MAFFT<sup>46</sup> | 7.520 | --auto; used for within- and between-cluster similarity only. |
| BLAST+<sup>49</sup> | 2.16.0 | blastp, E-value ≤ 10<sup>−5</sup>, word size 3, BLOSUM62, remaining parameters default; searched against human RefSeq<sup>48</sup> (downloaded Apr. 2026). |
| *Motif discovery and comparison* |  |  |
| HOMER<sup>31</sup> | 5.1 | findMotifsGenome.pl -size 200 on danRer11 (GRCz11); known motifs scored against JASPAR 2024 CORE vertebrates (redundant) at *p* ≤ 10<sup>−12</sup>, de novo motifs retained without a significance filter. compareMotifs.pl at defaults for motif clustering. |
| *Structure and statistical analysis* |  |  |
| Python | 3.12 | NumPy 2.4.6, SciPy 1.15.2, scikit-learn 1.5.2, pandas 2.2.3, ProDy 2.6.1. SciPy supplied the KD-tree used for side-chain–DNA contact extraction, the Mann–Whitney *U* test, and Pearson/Spearman correlations; scikit-learn supplied AUROC and average precision. Motif similarity implemented in NumPy. ProDy for mmCIF-to-PDB conversion. |
| ChimeraX<sup>50</sup> | 1.9 | Visualizing structures and calculating backbone RMSD of each predicted DBD to its closest PDB template. |
| *Downstream analysis* |  |  |
| chromVAR / motifmatchr<sup>51</sup> | 1.27.0 / 1.27.0 | R 4.4.1 / Bioconductor 3.20; BSgenome.Drerio.UCSC.danRer11 1.4.2. matchMotifs at p.cutoff = 5e-5; getBackgroundPeaks with 50 iterations; fixed seed before background sampling. |
| Signac<sup>52</sup> / Seurat<sup>53</sup> | 1.14.0 / 5.1.0 | Integrated multiome object; ATAC counts summed and RNA averaged within each SEACells metacell (∼75 cells; metacells of <20 cells dropped, 1,232 retained). |
| Cicero<sup>54</sup> | 1.3.9 (monocle3 branch) | Peak co-accessibility to associate accessible peaks with target genes in CellOracle GRNs. |
| CellOracle<sup>36</sup> | 0.18.0 (gimmemotifs 0.17.0) | Base-GRN scanning at FPR = 0.02, motif score threshold = 10, on danRer11; Bagging ridge regression *α* = 10; edges retained at *p* < 0.001 (per-edge *t*-test) and capped at the top 2,000 per cell type by absolute coefficient. |
| *Reference data* |  |  |
| CisBP<sup>10</sup> | 3.10 | Benchmark PWMs with PBM evidence; CisBP and Hybrid GRNs built using *D. rerio* PWMs. |
| JASPAR<sup>11</sup> | 2024, CORE vertebrates (redundant) | Ground-truth ortholog motifs, HOMER known-motif database, and the null distribution for empirical *p*-values. |
| AnimalTFDB<sup>30</sup> | 4.0 | 2,543 zebrafish TF sequences after removing chromatin remodelers and cofactors; family vocabulary for baselines. |
| ZFIN orthology<sup>47</sup> | Downloaded July 2026 | Curated zebrafish–human ortholog assignments. |
| Zebrahub-Multiome<sup>29</sup> | 2025 release | Paired snATAC + snRNA, 10–24 hpf; 640,834 peaks; precomputed Leiden peak clusters and SEACells partitions. |
| Genome | danRer11 (GRCz11) | Assembly for all motif scanning and peak coordinates. |

**Supplementary Video 1.**
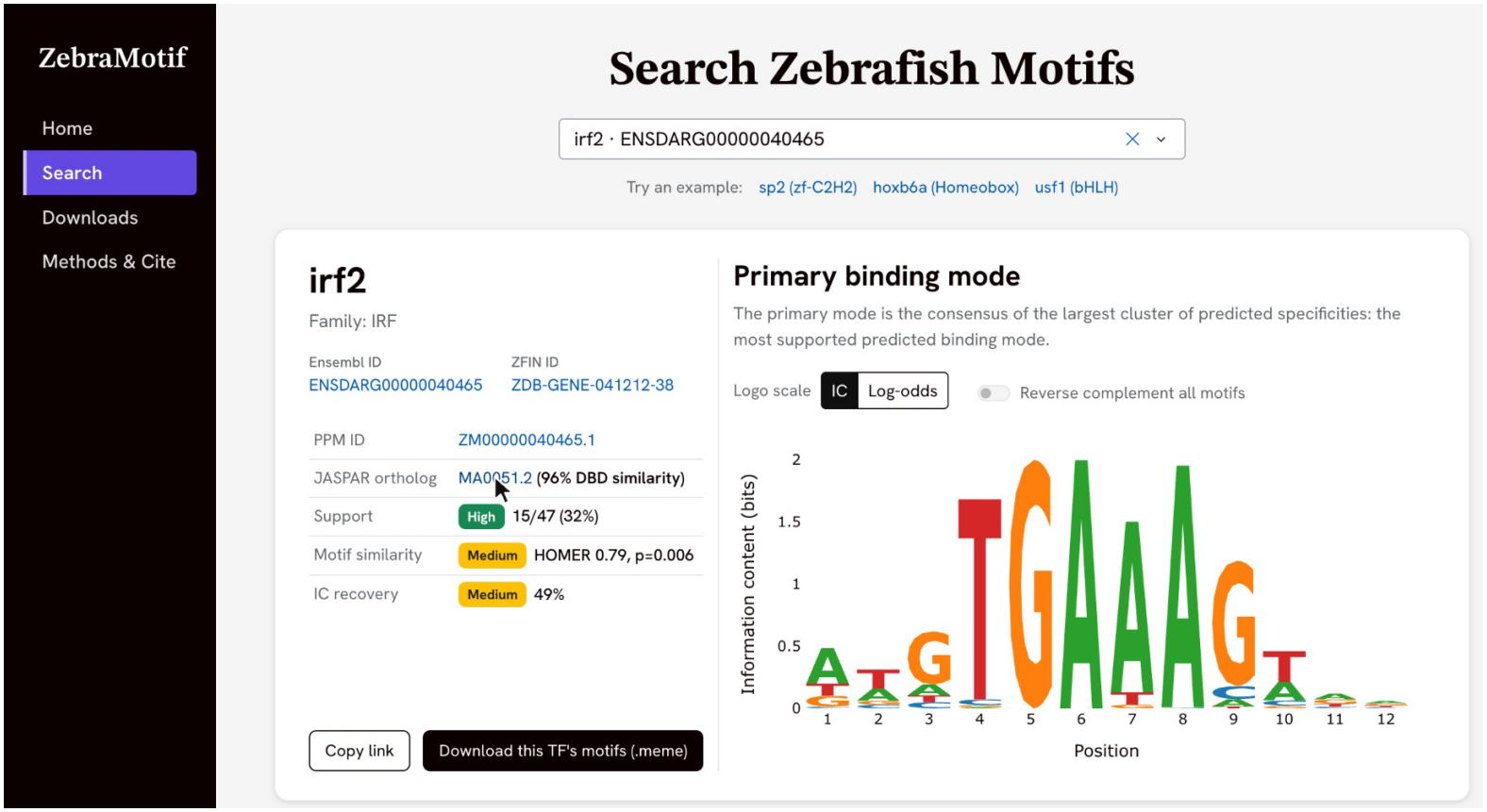
Walkthrough of the ZebraMotif website. Starting from the home page, the transcription factor *irf2* is found through the search field and its entry opens. The entry shows the TF’s identifiers (Ensembl, ZFIN), its closest JASPAR ortholog with DBD similarity, and the support, motif similarity and IC recovery of its primary binding mode, whose logo is then explored position by position (information content and base frequencies on hover) and switched between information-content and log-odds scales. Scrolling down shows the three alternate binding modes, aligned to the primary mode, and the predicted AF3 structure, which is rotated and zoomed, first restricted to the DNA-binding domain and then with the residues coloured by base or backbone contact with the DNA, next to its DBD-pLDDT and ipTM confidence. Screen recording of zebramotif.apps.czbiohub.org; 32 s, silent. The still above is at 7 s. Available at Zenodo (doi.org/10.5281/zenodo.23380219); file zebramotif_website_walkthrough.mp4.

